# Cross-trait assortative mating is widespread and inflates genetic correlation estimates

**DOI:** 10.1101/2022.03.21.485215

**Authors:** Richard Border, Georgios Athanasiadis, Alfonso Buil, Andrew Schork, Na Cai, Alexander Young, Thomas Werge, Jonathan Flint, Kenneth Kendler, Sriram Sankararaman, Andy Dahl, Noah Zaitlen

## Abstract

The observation of genetic correlations between disparate traits has been interpreted as evidence of widespread pleiotropy, altered theories of human genetic architecture, and spurred considerable research activity across the natural and social sciences. Here, we introduce cross-trait assortative mating (xAM) as an alternative explanation for observed genetic correlations. We observe that xAM is common across a broad array of phenotypes and that phenotypic cross-mate correlation estimates are strongly associated with genetic correlation estimates (*R*^2^ = 76%). Then, we present theoretical and simulation-based results demonstrating that, under xAM, genetic correlation estimators yield significant estimates even for traits with entirely distinct genetic bases. We demonstrate that existing xAM plausibly accounts for substantial fractions of genetic correlation estimates in two large samples (*N* = 827,960). For example, previously reported genetic correlation estimates between many pairs of psychiatric disorders are fully consistent with xAM alone. Finally, we provide evidence for a history of xAM at the genetic level using a novel approach based on cross-trait even/odd chromosome polygenic score correlations. Together, our results demonstrate that previous reports have likely overestimated the true genetic similarity between many phenotypes.

## 1 Introduction

Over the previous decade, considerable research activity has focused on using genetic data to infer the extent to which pairs of complex traits share overlapping genetic bases [1–7]. These efforts have been propelled by the development of genetic correlation estimators designed to use summary statistics from genome-wide association studies (GWAS) [3, 8, 9], which have become a fundamental statistical tool across many domains of human complex trait genetics, including psychiatry [2, 10], economics [11, 12], and medicine [13, 14]. The results of these analyses have been striking: many trait pairs, even those with limited phenotypic similarity, display nontrivial genetic correlations (e.g., 0.209 [*se* = 0.042] for Attention-deficit Hyperactivity Disorder [ADHD] and body mass index [BMI] in [2]). These findings have been broadly interpreted as evidence for widespread pleiotropy across the phenome [3–5, 15], and, in the case of psychiatric disorders, have raised concerns about the suitability of the existing nosology in the face of shared pathogenesis at the genetic level across ostensibly distinct phenotypes [2, 7].

However, such interpretations assume that genetic correlation estimators index shared genetic architectures, and various alternative explanations have been explored in considerable detail (e.g., vertical pleiotropy and diagnostic errors [5, 16]). Here, we consider a previously unexamined source of potential bias: cross-trait assortative mating (xAM), the phenomenon whereby mates are correlated across distinct traits (i.e., individuals’ values of phenotype *Y* are correlated with their partners’ values of phenotype *Z*). There are at least four reasons to be concerned with this potential oversight: First, the single trait random effects model, which genetic correlation estimators generalize, is misspecified under single-trait AM and as a result overestimates SNP heritability [17]. Second, single-trait AM is widespread across multiple domains for which substantial genetic correlations have been observed, including anthropometric, psychosocial, and disease traits [2, 4, 5]. Third, recent work has provided genetic-level evidence for a history of single-trait AM with respect to some of these same phenotypes [18]. Fourth, xAM is known to generate spurious results for other marker-based inference procedures, including Mendelian randomization [19] and GWAS [20].

The present investigation comprises the first systematic investigation of the impact of xAM on genetic correlation estimates. We first compile the largest to-date atlas of cross-mate cross-trait phenotypic correlations, examining a broad array of previously-studied metabolic, anthropometric, psychosocial, and psychiatric phenotypes across two large population based samples in the United Kingdom (*n*=81,394) and Denmark (*n*=746,566). We find that these cross-mate correlations, which are derived from phenotypic measurements, explain a major portion of empirical marker-based genetic correlation estimates for the same trait pairs (*R*^2^=76% across datasets). We next present theoretical and simulation-based results demonstrating that xAM biases genetic correlation estimates and yields significant estimates even among traits with uncorrelated genetic effects. We then use a simulation-based approach to evaluate the extent to which empirical level of xAM alone might plausibly explain empirical genetic correlation estimates among previously studied traits, finding that substantial fractions of empirical genetic correlation estimates are congruent with expectations for biologically independent traits subject to xAM. Lastly, we present a multivariate extension of the work of Yengo and colleagues [18], who utilized correlations between even versus odd chromosome-specific polygenic scores to detect genetic signatures of single trait AM. We find that cross-trait even/odd polygenic score correlations mirror cross-mate phenotypic correlation patterns and, through this association, can explain substantial variation in empirical genetic correlation estimates.

Together, our findings imply that xAM-induced artifacts comprise a common, previously unaddressed systematic source of bias in the genetic correlation literature. We discuss these results in the context of the widespread application of the genetic correlation as a measure of shared etiology, suggesting that many previously reported genetic correlations are overestimated.

## 2 Results

### 2.1 Genetic correlation estimates mirror cross-mate phenotypic correlations

We begin by quantifying the extent to which empirical genetic correlation estimates align with cross-trait spousal correlations across a broad array of previously studied phenotypes: a set of 20 anthropometric, metabolic, and psychosocial traits measured in the UK Biobank (UKB) [21] and a collection of five psychiatric disorder diagnoses ascertained from Danish civil registry data [22–26]. We first i dentified 40,697 sp ousal pa irs within the UKB sample and randomly selected 373,283 mate pairs within the Danish sample and then estimated cross-mate correlations. For a pair of phenotypes *Y*, *Z*, there are three cross-mate correlation parameters: *r*_*yy*_ (resp. *r*_*zz*_) the correlation between mates on phenotype *Y* (resp. phenotype *Z*) and *r*_*yz*_, the cross-mate cross-trait correlation; we generically denote these quantities *r*_mate_, and present these estimates in the diagonal and sub-diagonal entries of Figures 1a and 1b. We also compiled LDSC genetic correlation estimates, denoted 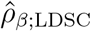, for each pair of phenotypes, which we present in the super-diagonal entries of Figures 1a and 1b. In the context of an ordinary least squares (OLS) regression model, 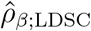 were strongly associated with *r*_mate_ estimates across both sam-ples (Figure 1c; meta-analytic *R*^2^=75.99%, 95% CI: 68.60% - 83.39%). We also fit a Bayesian linear model accounting for heteroskedasticity and sampling error in *r*_mate_ estimates, which yielded comparable results (*R*^2^=76.95%, 95% CI: 74.19% - 79.72%). All pairwise estimates are provided in Table S1.

**Figure 1:**
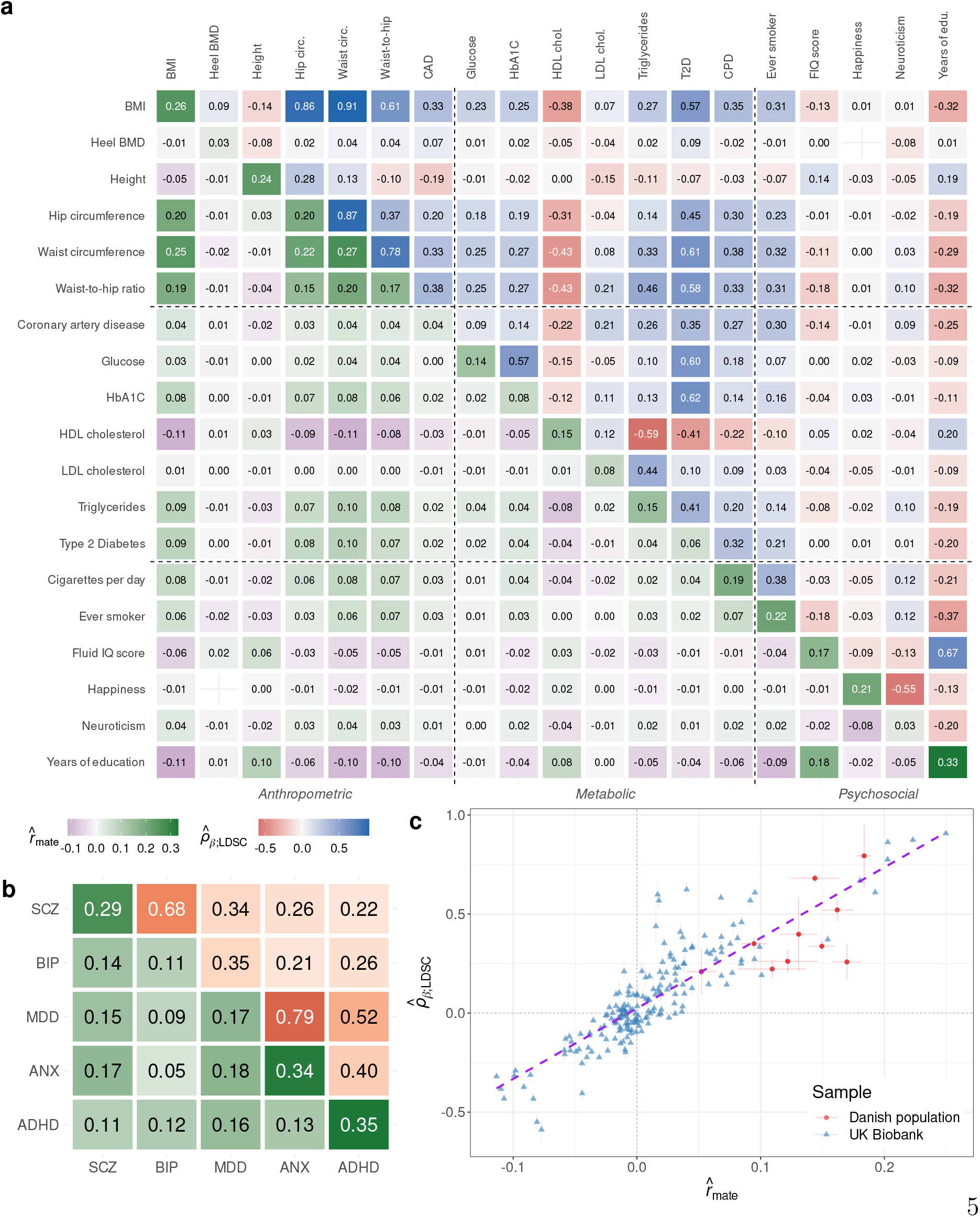
(a) Cross-mate correlation and genetic correlation estimates for previously studied UK Biobank phenotypes. Diagonal and sub-diagonal heatmap entries correspond to cross-mate phenotype correlation estimates derived from 40,697 putative spouse pairs in the UK Biobank. Super-diagonal entries correspond to empirical LDSC correlation estimates among unrelated European ancestry UK Biobank participants. Phenotypic correlations are sex- and age-adjusted partial correlation estimates. (b) Cross-mate correlation and genetic correlation estimates for psychiatric disorders. Diagonal and sub-diagonal entries reflect cross-mate tetrachoric correlations among 373,283 spousal pairs sampled from the Danish population. Super-diagonal entries are previously-reported LDSC correlation estimates [2]. (c) Association between empirical cross-mate phenotypic correlation and genetic correlation estimates (meta-analytic *R*^2^ ≈ 76%). Note: *all numbers have been rounded to two decimal places; the sex-constrained structural model for the cross-mate correlation between heel bone mineral density (BMD) and subjective happiness failed to converge and is omitted from the first heatmap*.

### 2.2 Defining genetic correlation

Having established that a large degree of the variance in genetic correlation estimates can be predicted from phenotypic mating correlations, we now provide theoretical intuition as to why this might occur. We start by defining two distinct notions of genetic similarity between phenotypes.

We consider a pair of phenotypes *Y*, *Z* composed of the additive effects of *m* standardized haploid variants *X*_1_, … *X*_*m*_ with phenotype-specific effects *β*_*y*_, *β*_*z*_:

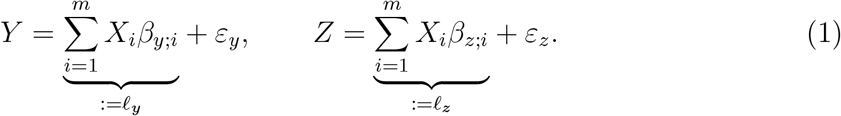

Each phenotype is composed of a heritable liability component *l*, the true polygenic score, and a non-heritable component *ε*. For convenience, we assume that causal variants are initially unlinked and both *Y* and *Z* have unit variance under random mating (panmixis), such that the panmictic heritabilities are 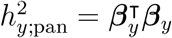 and 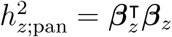.

Classically, genetic correlation is defined as the correlation between the heritable components of two traits [5, 27]. We will refer to this quantity as the score correlation:

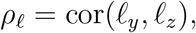

as it reflects the correlation between the true polygenic scores. This is distinct from the correlation between effects, which we refer to as the **effect correlation** *ρ*_*β*_, and which indexes the similarity of variant effects on two phenotypes:

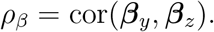

We will refer to a pair of traits as *genetically orthogonal* when their effects are uncorrelated; i.e., when *ρ*_*β*_ = 0. Variance components models aim to estimate *ρ*_*β*_ [28–30]. Within the standard random effects model framework, which assumes genotypes are independent of effect sizes, *ρ*_*l*_ and *ρ*_*β*_ are equivalent and hence seldom discussed as separate quantities.

We note that both of these measures are distinct from *pleiotropy*, which refers to the extent to which two traits are affected by the same causal variants. Pleiotropy is a necessary precondition for effect correlation, but it is possible to have independent effects with o4r without pleiotropy (Section 5.1.2).

Less intuitively, score correlations (*ρ*_*l*_) can be nonzero when the effects themselves are independent (i.e., *ρ*_*β*_=0). That is, traits with uncorrelated effects can have collinear polygenic scores. Under xAM, all causal variants affecting trait *Y* become correlated with all causal variants affecting trait *Z* and these correlations are directionally consistent with their respective effects (see Supplementary Materials S1.1 for details). For example, conditional on having inherited a trait-increasing allele for phenotype *Y* from one parent, an individual is more (resp. less) likely to have inherited a trait-increasing (resp. - decreasing) allele for phenotype *Z* from the other parent when parents have positively assorted across *Y* and *Z*. As we will demonstrate in the following section, this results in a non-zero genetic correlation *ρ*_*l*_ in the direction of the cross-mate cross-trait phenotypic correlation, even for genetically orthogonal traits.

### 2.3 The impact of xAM in simulations

We ran a series of forward-time simulations using realistic genotype data (see Online Methods) to investigate the impact of xAM on multiple measures of genetic correlation. We began with a founder population of 335,550 of unrelated European ancestry UK Biobank participants’ phased haploid genotypes at 1,000,000 common single nucleotide polymorphisms (SNPs). We then simulated two phenotypes, *Y* and *Z*, by randomly selecting 10,000 causal variants and drawing two corresponding sets of standardized effects from the bivariate Gaussian distribution with variances 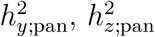 and correlation *ρ*_*β*_. Non-heritable components *ε*_*y*_, *ε*_*z*_ were drawn independently from univariate Gaussian distributions with variances 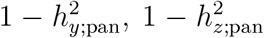, such that both phenotypes had unit variance at panmixis.

At each generation, individuals (consisting of a set of genotypes together with two phenotypes) were matched to achieve target cross-mate correlation parameters. Throughout this section, we assume the cross-mate phenotypic correlations are non-negative and exchangeable for simplicity; i.e., that *r*_*yy*_, *r*_*zz*_, and *r*_*yz*_ are all equal to a single parameter, which we denote *r*_mate_ ≥ 0. Extension to dissassortative mating, wherein *r*_mate_ < 0, is trivially achieved by reversing the sign of either phenotype. We also assume the two phenotypes have equal panmictic heritabilities, which we denote 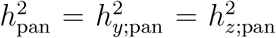. We later consider general cross-mate correlation structures and differing panmictic heritabilities.

We constructed offspring genotypes by simulating meiosis according to empirically derived recombination frequencies in order to preserve local LD. Each generation’s haploid genotypes were thus permutations of the founder haplotypes. We performed sensitivity analyses (see Online Methods) to confirm that our results did not depend on simulation parameters, including the number of causal variants (Figure S1), mate selection algorithm (Figure S2), recombination scheme (Figure S3), and whether causal variants with orthogonal genetic effects were located on overlapping loci (Figure S4). Having constructed new genotypes, we generated phenotypes as above and estimated effect correlation 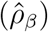 using LD score regression (LDSC, denoted 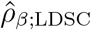; [3]), Haseman-Elston regression (HE, denoted 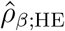; [31]), and residual maximum likelihood (REML; 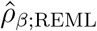; [28, 32]). We also computed true score correlations (*ρ*_*l*_), which is possible when the true genetic effects are known. Additionally, we conducted supplementary simulation studies investigating the impact of xAM on GWAS effect size estimates (Section 5.6, Figure S5) and the impact of xAM on effect correlation estimates in the presence of misdiagnosis (Supplementary Materials S1.2.2, Figure S6).

#### 2.3.1 xAM induces nonzero score correlations among genetically orthogonal traits

We confirmed that xAM induces substantial score correlations across abroad array of simulation parameters. This was perhaps most striking for traits with orthogonal effects: Figure 2a demonstrates the increase in the true score correlation across multiple generations of xAM for a pair of traits with *ρ*_*β*_ = 0, *r*_mate_ = 0.5, and 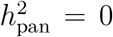. Averaging across simulation replicates, after a single generation of xAM, the score correlation was 0.11, and after three generations of xAM, the score correlation was 0.24. The fraction of shared causal variants had no discernible impact on this behavior (Figure S4); that is, when *ρ*_*β*_ = 0, the extent to which the causal variants affecting two traits overlapped was unimportant.

**Figure 2:**
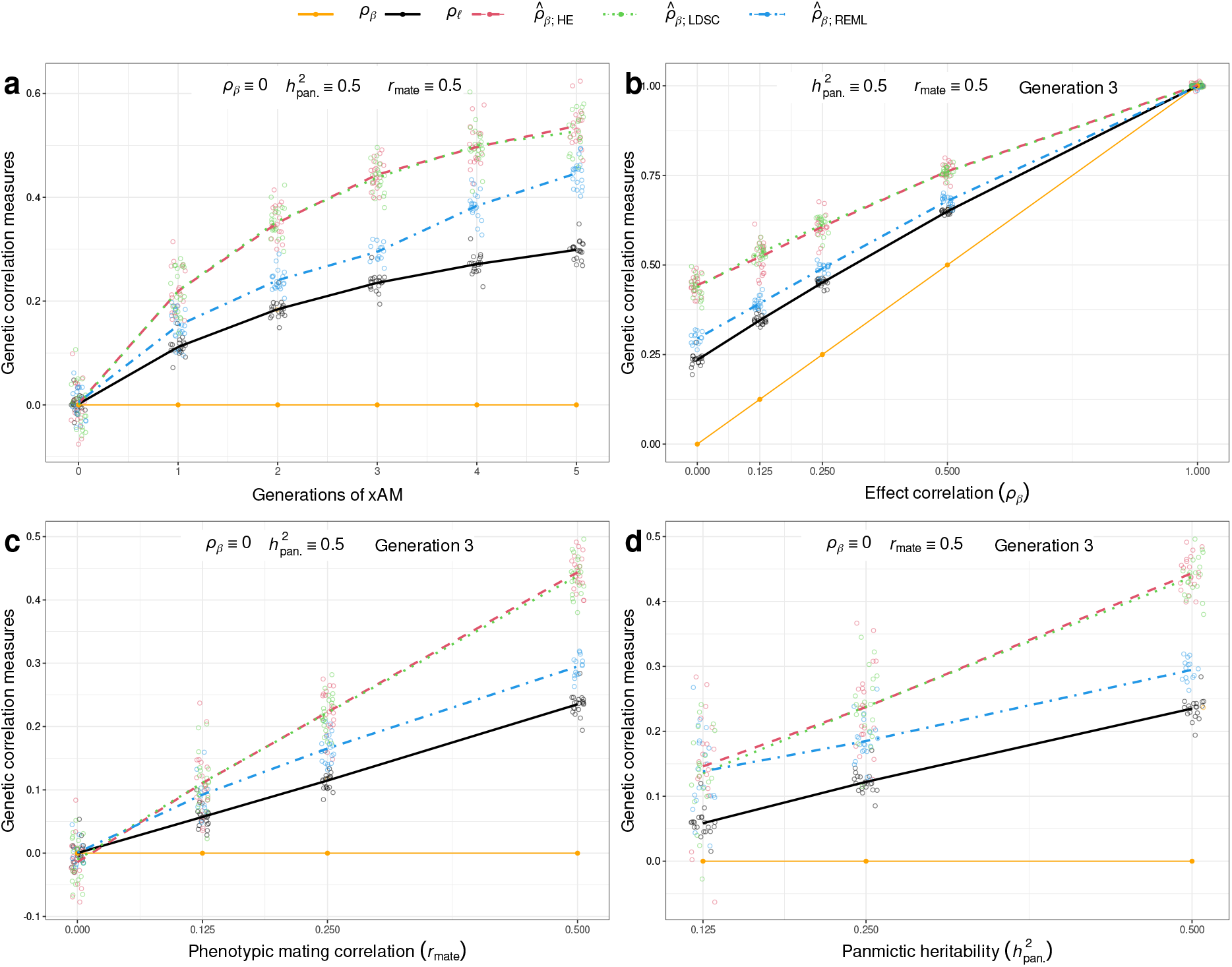
Forward-time simulation results demonstrating the impact of xAM on score correlation (*ρ*_*l*_) and effect correlation estimates 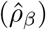 for two synthetic phenotypes with true effect correlation *ρ*_*β*_ and panmictic heritabilities 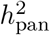. The single mating correlation parameter *r*_mate_ denotes the value of the two cross-mate single-trait correlations and the cross-mate cross-trait correlation. (a) xAM increases the true score correlation among two genetically orthogonal phenotypes. The HE, LDSC, and REML estimators all overestimate *ρ*_*β*_, and the magnitude of this bias increases over subsequent generations. (b) After three generations of xAM, 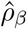 estimates are upwardly biased for trait pairs with true effect correlations less than one (i.e., genetically distinct phenotypes). (c) The impact of three generations of xAM increases with the phenotypic correlation among mates. (d) The impact of three generations of xAM increases monotonically with the panmictic heritabilities.

Intuitively, this behavior occurs because xAM induces correlations between causal loci consistent in sign with the directions of their effects (see S1.1.1 for details). That is, pairs of trait increasing or trait decreasing loci become positively correlated whereas pairs of loci discordant with respect to the direction of their effects become negatively correlated. We can decompose the liability correlation into two disjoint sums:

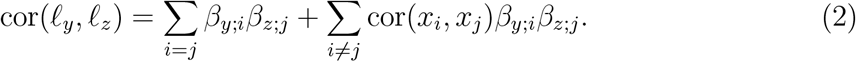

The first sum is proportional to *ρ*_*β*_ and will only be nonzero when *ρ*_*β*_ ≠ 0. On the other hand, the second sum does not depend on *ρ*_*β*_, but, due to sign-consistent global LD induced by xAM, will be greater than zero at all pairs of causal loci for *Y* and/or *Z*.

#### 2.3.2 Effect correlation estimates are biased upwards

Although xAM induces nonzero score correlations between genetically orthogonal traits, these correlations are not artifactual in the sense that individuals’ latent liabilities truly do become correlated. On the other hand, effect correlation estimators, which assume that the contributions of causal haplotypes are pairwise independent (i.e., cov(*X*_*i*_*β*_*y*;*i*_, *X*_*j*_*β*_*z*;*i*_) = 0 for distinct loci *i*, *j*) [17], are mispecified under xAM and yield upwardly biased estimates of the true effect correlation *ρ*_*β*_. Returning to the case of genetically orthogonal traits presented in Figure 2a, after a single generation of assortative mating, the REML estimator yielded 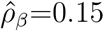 and the method-of-moments estimators, HE and LDSC, which are mathematically equivalent to each other [33, 34], yielded average estimates of 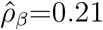 and 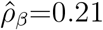, respectively, all of which are substantally higher than the true effect correlation of zero. After three generations of xAM, this upward bias becomes more pronounced, with REML and LDSC yielding effect correlation estimates of 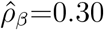 and 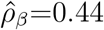, respectively.

#### 2.3.3 The true score correlation and estimated effect correlation are monotonically related to *ρ*_*β*_, *r*_mate_, and 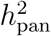

So far we have our focus to the instructive case of genetically orthogonal traits, though many trait pairs subject to cross-trait xAM will truly have correlated genetic effects. Figure 2b illustrates the relationship between *ρ*_*β*_, *ρ*_*l*_, and 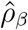 for two traits with 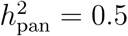 and subject to three generations of xAM with *r*_mate_ = 0.5. Excepting the case of *ρ*_*β*_ = 1.0 (a pair of genetically identical phenotypes), the pattern of results remains consistent with the genetically orthogonal case, where the correlation between effects is lower than the true score correlation, which is in turn lower than the upwardly biased 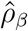 estimates provided by REML, HE, and LDSC. For example, when *ρ*_*β*_ = 0.25, LDSC yields 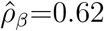 after three generations of xAM. The impact of cross-trait xAM on both *ρ*_*l*_ and 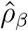 is greater for traits subject to higher cross-mate correlations (i.e., stronger assortment induces more substantial bias; Figure 2c) and for traits with greater panmictic heritabilites (i.e., the genetic consequences of xAM are more substantial for more heritable phenotypes; Figure 2d).

### 2.4 xAM alone can plausibly explain substantial empirical genetic correlation estimates

Having established in simulations that xAM can induce substantial 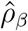 estimates regardless of shared biological basis, we sought to quantify the extent to which empirical 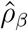 estimates for previously-studied trait pairs could be explained by xAM alone, assuming genetic orthogonality. As there are currently no unbiased estimators of *ρ*_*β*_ under xAM, we proceeded with a simulation-based approach, restricting our focus to method-of-moments estimators in light of the widespread usage of LDSC regression. We present two self-contained analyses focused on the UKB (Figure 1a) and Danish sample phenotypes (Figure 1b), respectively.

Within each sample, we first identified mate pairs and estimated the phenotypic cross-mate correlations *r*_*yy*_, *r*_*zz*_, and *r*_*yz*_ for each pair of traits as described above. We then used these estimates, together with empirical heritability estimates, as inputs to a forward-time simulation where separate, non-overlapping collections of causal variants were assigned to the two phenotypes, thereby constraining the true effect correlation to zero. At each generation, we estimated the effect correlation, *ρ*_*β*_, using method-of-moments. These projected effect correlation estimates, which we denote 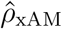, can be interpreted as the expected LDSC effect correlation estimate for the two traits in the complete absence of pleiotropy after a given number of generations of xAM with mating proceeding in accordance with empirical spousal correlation estimates.

Given that both marker-based and pedigree-based heritability estimators (which we denote 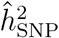 and 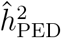, respectively) are biased under AM (albeit in different directions;[17, 35]) and that neither approach directly estimates the panmictic heritability, we conducted two sets of simulations, one assuming 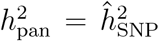 and the other assuming 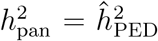. We focus discussion in the primary manuscript on the simulations which assumed 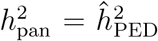 as the impact of xAM depends on the total heritability of the traits involved, not just that tagged by common variants, though results assuming 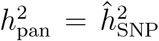 are presented in the supplement.

We next compared the projected correlation estimates under xAM alone, 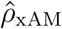 to empirical LDSC effect correlation estimates using real (rather than synthetic) phenotype data, which we denote 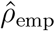 To simplify discussion, we define the ratio

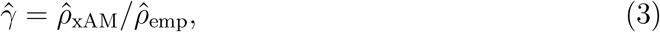

which measures the strength of the projected LDSC effect correlation estimate due to xAM-induced artifact relative to the empirical LDSC effect correlation estimate for a given pair of phenotypes. When 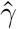 close to zero, empirical effect correlation estimates are far greater than what might be attributable to xAM alone. On the other hand, when 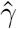 is close to one, empirical estimates are entirely consistent with xAM-induced artifact. We caution that 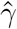 does not constitute a direct estimate of the fraction of the empirical genetic correlation estimate attributable to xAM.

#### 2.4.1 Expected effect correlation estimates for UK Biobank phenotypes in the absence of pleiotropy

As we were concerned with the misinterpretation of xAM-induced artifacts for evidence of pleiotropy, we restricted our attention to 132 (of 190 possible) pairs of phenotypes with nominally significant LDSC genetic correlation estimates (Table S1). We first obtained pedigree-based heritability estimates for each of the traits of interest from the existing literature, using estimates derived in UK-based samples when available (Table S2). Together with the phenotypic mating correlations (Figure 1a), these comprised inputs for the forward time simulations used to compute 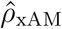, which we in turn compared to empirical LDSC effect correlation estimates 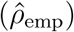 to obtain 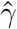 as illustrated in Figure 3a.

**Figure 3:**
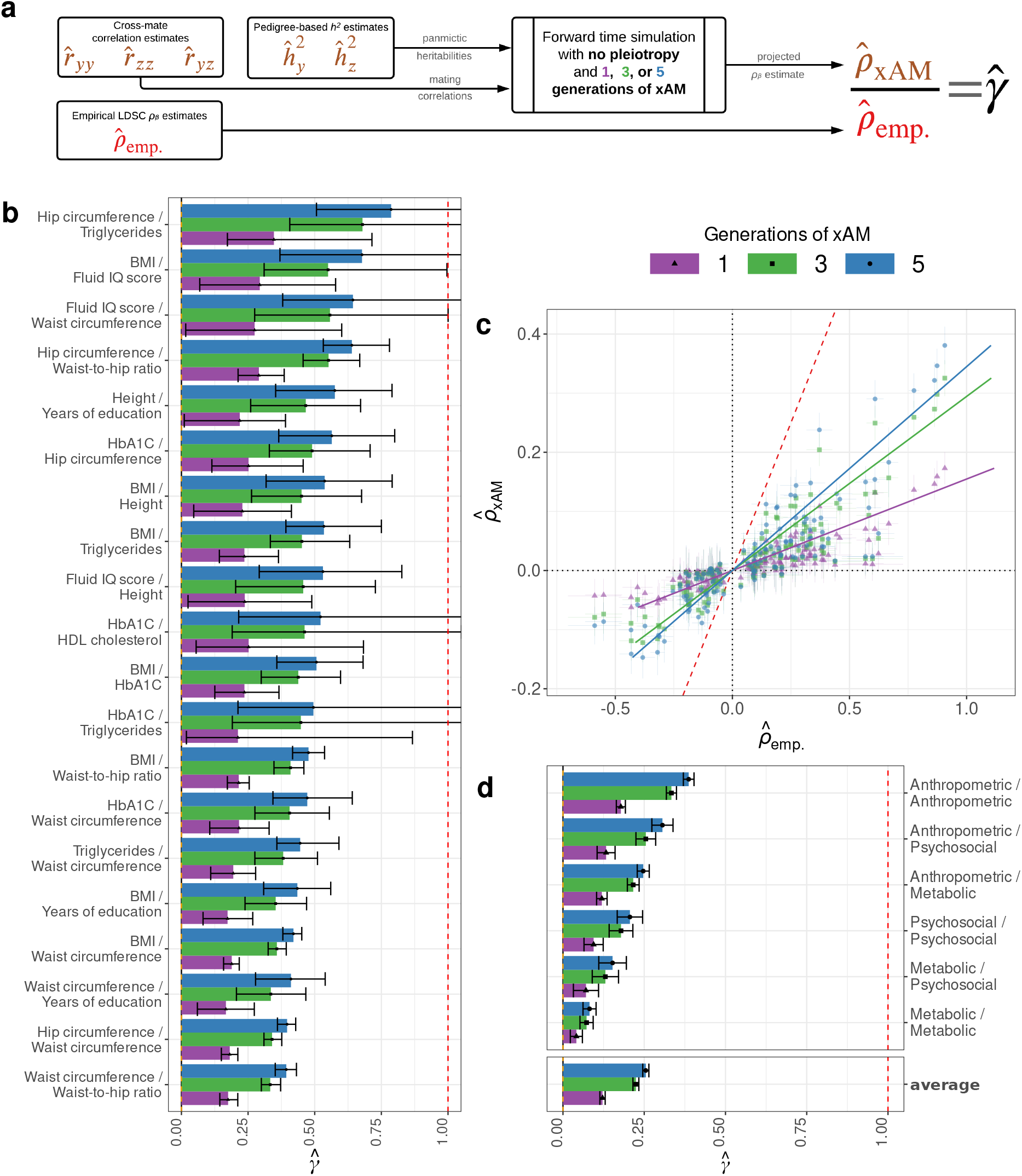
Comparison of empirical genetic correlations 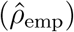 to expected estimates in the absence of pleiotropy 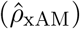 for assorted UK Biobank phenotypes. Error bars throughout represent 95% credible intervals. (a) For each trait pair, we ran 400 replicate forward time simulations without pleiotropy with panmictic heritabilities obtained from pedigree-based estimates and selecting mates to match empirical cross-mate correlation estimates. After each generation, we computed the method-of-moments genetic correlation estimate 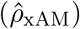, which we interpret as the expected LDSC genetic correlation estimate in the absence of pleiotropy after a given number of generations of xAM. We compared this quantity to empirical LDSC genetic correlation estimates in the UK Biobank 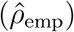 to obtain the ratio 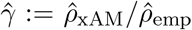. (b) The top 20 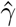 estimates after up to five generations of simulated xAM across previously-studied UK Biobank phenotype pairs with nominally significant LDSC genetic correlations. (c) Projected versus empirical LDSC estimates for all UK Biobank phenotypes with nominally significant genetic correlation estimates. (b) Inverse variance weighted average 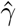 estimates within and between UK Biobank phenotypes in qualitatively similar domains.

Across 132 trait pairs, 42 evidenced 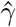 values significantly greater than zero after a single generation of xAM, which increased to 74 trait pairs after three generations. Figure 3b presents the first 20 pairs in descending order of 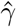 and Figure 3c presents the raw projected and empirical effect correlation estimates across all 132 pairs (see Table S3, Figures S7 to S9 for detailed results spanning one to five generations of xAM). Averaging across all trait pairs (including those not significantly different from zero), the inverse variance weighted average 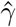 estimate was 0.25 (*se*=0.005). The relative strengths of within-and cross-domain projections varied substantially when aggregating across qualitative clusters of phenotypes (see Figure 1a and Table S2 for domain definitions). For example, relatively small fractions of empirical 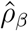 estimates among metabolic phenotypes were congruent with expectations under five generations of xAM alone 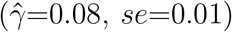 in comparison to those between psychosocial phenotypes 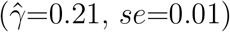. We note that these domain-averages are highly dependent on the set of traits considered and should be interpreted with care.

#### 2.4.2 Expected effect correlation estimates among psychiatric disorders in the absence of pleiotropy

We next computed projected effect correlation estimates assuming up to five generations of xAM under empirically derived cross-mate correlation structures for a collection of five psychiatric disorders, as estimated in sample of 373,283 spousal pairs randomly selected from the Danish population (see S1.3 for details). Again, we obtained pedigree-based heritabilities for each disorder, preferring studies using samples from Scandinavian countries when available (Table S4), and used these estimates, together with the cross-mate-correlations, to compute 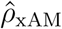. We then compared these projections to the LDSC genetic correlation estimates presented by Anttila and colleagues [2]. We focused on Anttila et al.’s manuscript as their interpretation of the genetic correlation as a direct measure of biological overlap is broadly representative of the views of many complex trait geneticists [3–5, 7].

Across all pairwise combinations of disorders, we observed an average ratio of 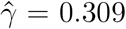 (*se*=0.037; Figures 4a and 4b) after five generations of x AM. Some trait p airs evidenced considerably greater empirical effect correlation estimates than might be explained by xAM alone (e.g., for anxiety disorders and major depression, 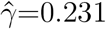, 95% CI: 0.129 −0.390), whereas for other pairs this discrepancy was modest (e.g., for anxiety disorders and Schizophrenia, 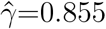, 95% CI: 0.461 - 2.583). We present detailed results across one to five generations of xAM in Table S5 and Figures S10 and S11.

**Figure 4:**
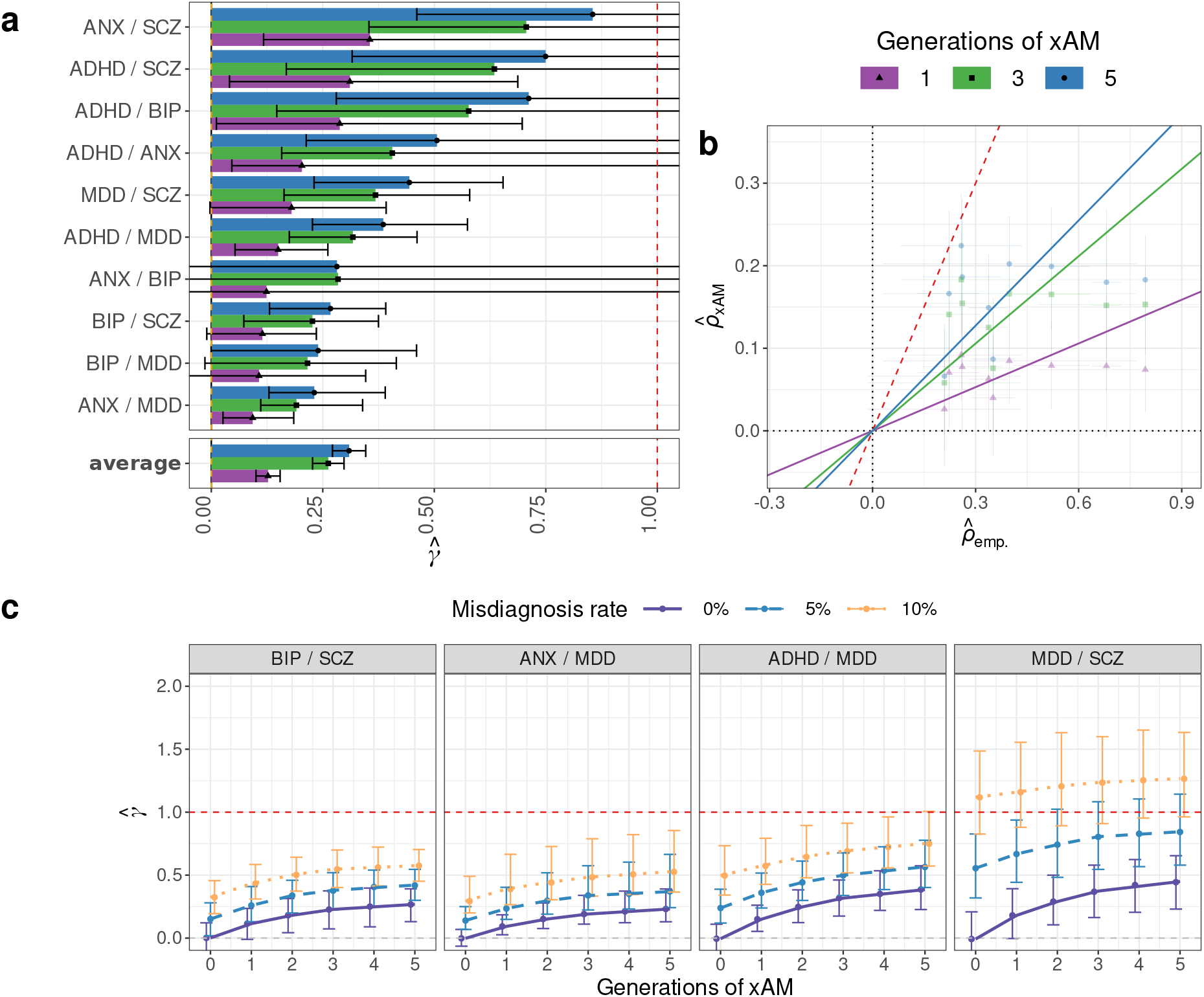
Comparison of empirical effect correlation estimates among psychiatric phenotypes to expectations in the absence of pleiotropy. (a) Ratios 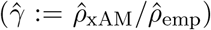 of projected LDSC effect correlation estimate 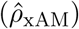 assuming xAM and no pleiotropy relative to the empirical effect correlation estimates 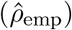 of [2] for five psychiatric disorders with 95% credible intervals. Projections assume up to five generations of xAM matching empirical cross-mate correlations in the Danish sample and assume panmictic heritabilities equal to pedigree-based estimates from demographically-comparable studies (Table S4). (b) Projected versus empirical LDSC estimates across psychiatric phenotype paits. (c) The potential joint impacts of bidirectional errors in diagnosis and xAM on effect correlation estimates for selected psychiatric disorder pairs. The red dashed line corresponds to 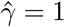 across all panes.

##### xAM exacerbates the impact of misdiagnosis on effect correlation estimates among psychiatric disorders

Given the combined effects of xAM and measurement error in terms of inflating effect c orrelation estimates (see S 1.2.2), we next extended our method for deriving 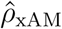 to incorporate errors in diagnosis and applied this extension to each pair of psychiatric disorders. At each generation, we dichotomized continuous phenotypes using empirical prevalence estimates (Table S4) and introducing misdiagnosis errors prior to estimating the effect correlation. We considered bidirectional misclassification errors (one disorder might be mistaken for another and vice-versa) ranging from 0% to 10%and up to three generations of xAM in accordance with empirical spousal-correlation estimates (Figure S11). Results demonstrated that some of the empirical effect correlation estimates presented by [2] were entirely consistent with artifacts induced by xAM coupled with moderate diagnostic error rates for some trait pairs but not others. Whereas high (i.e., ≥10%) rates of misdiagnostic errors are necessary to fully account observed effect correlations between ADHD and anxiety disorders in the absence of xAM, five generations of xAM coupled with a 5% misdiagnosis rate would account for nearly all of the empirical genetic correlation estimate (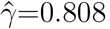, 95% CI: 0.363 - 5.106). On the other hand, the same extent of xAM and misdiagnosis would account for a considerably smaller fraction of the empirical genetic correlation estimate between anxiety disorders and major depression (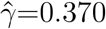, 95% CI: 0.242 − 0.665). Figure 4c highlights the potential impacts of xAM and diagnostic errors on four selected trait pairs.

### 2.5 Genetic evidence for xAM recapitulates empirical cross-mate correlations

We have thus far established that one or more generations of xAM consistent with empirical cross-mate correlation estimates could plausibly induce substantial bias in genetic correlation estimates for many previously studied traits. We now assess evidence for a history of phenotypically-mediated xAM at the genetic level. Previous work has used even/odd chromosome polygenic score correlations to examine evidence for AM with respect to a single trait [18]. We adapted this approach to the bivariate case by computing *cross-trait* even/odd polygenic score correlations (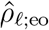; Table S6). In the absence of xAM or other sources of population structure, polygenic scores computed on even chromsomes for one trait versus those computed on odd chromosomes for a second trait are expected to be orthogonal. On the other hand, the long-range sign-consistent LD induced by xAM will induce correlations between polygenic scores on separate chromosomes (see Supplemental Materials S1.1.3). Further, these correlations should positively associate with cross-mate phenotypic correlations.

In order to compute cross-chromosome polygenic score correlations, we first computed GWAS effect estimates and clumped results in a training subsample of 80% of unrelated European ancestry UK Biobank participants and then computed polygenic scores specific to even and odd chromosomes for the remaining 20% using multiple *p*-value thresholds. We then regressed the estimated even/odd polygenic score correlations on the corresponding cross-mate cross-trait correlations (or cross-mate same-trait correlations in the case of univariate even/odd correlations; Section 5.9). Finally, we regressed the estimated cross-trait even/odd polygenic score correlations on the corresponding LDSC effect-correlation estimates.

In an OLS model using scores built with all clumped variants, cross-mate phenotypic correlation estimates (*r*_mate_) explained substantial variance in the cross-chromosome even/odd polygenic score correlations (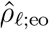; *R*^2^=47.66%, *se*=0.060; Figure 5a). As before, accounting for measurement error and heteroskedasticity in a Bayesian linear model yielded nearly identical results. This association persisted across polygenic score *p*-value thresholds ranging from 3.00e-3 to 1.00; Figure S12).

**Figure 5:**
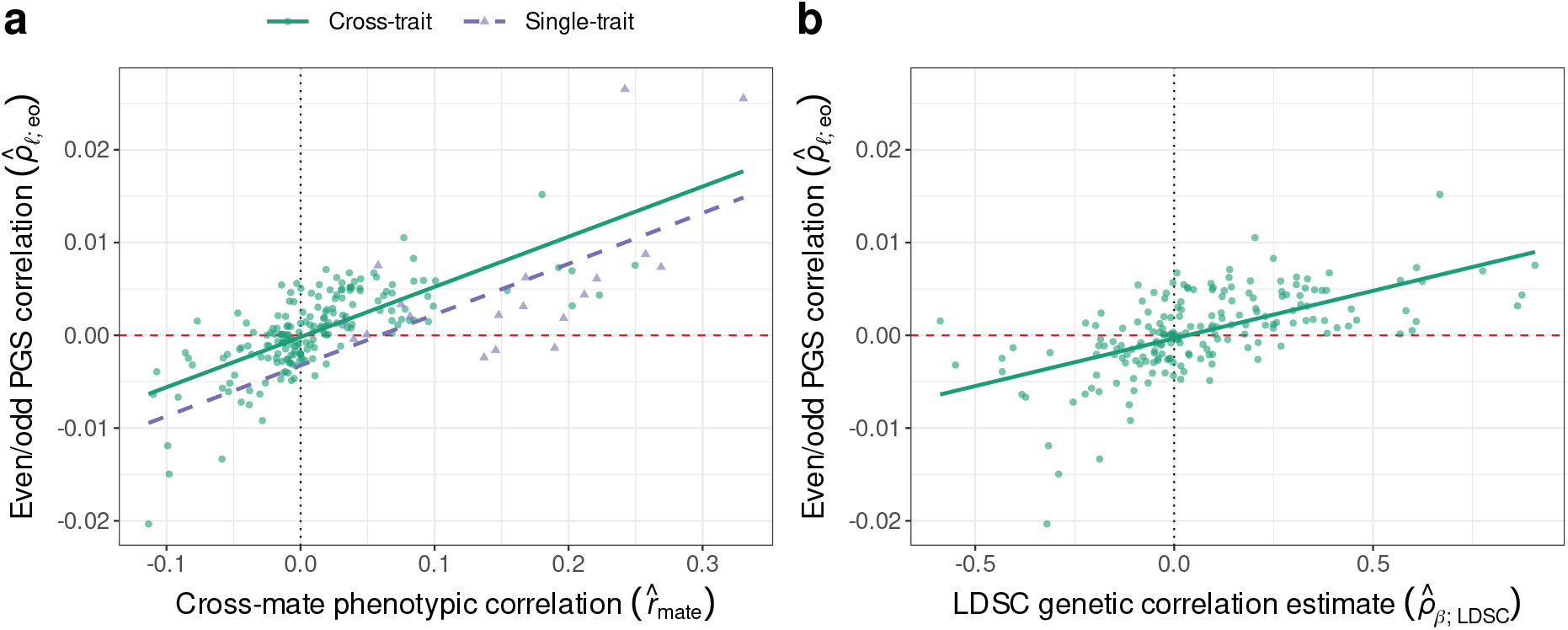
Genetic level evidence consistent with xAM in the UK Biobank. (a) Correlation between even and odd chromosome-specific polygenic scores as a function of the cross-mate phenotypic correlation. For a single trait, the vertical axis reflects the correlation between even and odd chromosome polygenic scores 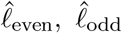 and the horizontal axis reflects the cross-mate correlation. For a pair of traits *Y*, *Z*, the vertical axis reflects a single parameter to which the correlations between 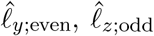 and between 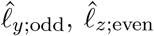 are both constrained, and the horizontal axis reflects the cross-mate cross-trait correlation. (b) Cross-trait even/odd polygenic score correlations as a function of empirical LDSC effect correlation estimate. The positive association of these quantities is consistent with our hypothesis that empirical effect correlations capture genetic structure in addition to signals of biological overlap.

Additionally, cross-trait even/odd chromosome polygenic score correlations were positively associated with empirical LDSC effect correlation estimates (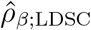; *R*^2^=34.81%, *se*=0.045; Figure 5b). This is consistent with the hypothesis that empirical effect correlation estimates are capturing additional structure beyond the signatures of biological overlap. Further, regressing 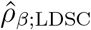 on 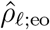 and 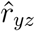 simultaneously revealed that the association between 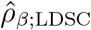 and 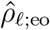 is mediated via 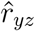 (Δ*R*^2^<0.001; partial effect *p*=0.48 for 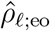 versus *p*<5e-8 for 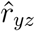). Thus, any alternative sources of population structure beyond xAM captured by 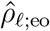 do not appear to explain the positive association between 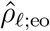 and 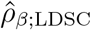.

## 3 Discussion

Nonzero effect correlation estimates have been widely interpreted as evidence that two traits share overlapping genetic bases. It is therefore surprising that substantial variation in genetic correlation estimates can be explained by cross-mate phenotypic correlations (Figure 1c). Further, given the strength of this association, the consequences of the random mating assumption implicit in all commonly used genetic correlation estimators [3, 8, 28, 31, 36] warrant critical attention.

We have demonstrated that xAM increases the score correlation (*ρ*_*l*_) between a pair of traits by inducing long-range sign-consistent LD across all pairs of causal variants, even when the causal effects themselves are uncorrelated (*ρ*_*β*_ = 0). Marker-based genetic correlation estimators, however, implicitly assume that *ρ*_*l*_ = *ρ*_*β*_, and under this misspecification substantially overestimate *ρ*_*β*_ after even a single generation of x AM (Figure 2a). In this context, we sought to quantify this potential source of bias across a broad array of phenotypes by compiling the largest to-date collection of empirical cross-mate cross-trait correlation estimates and using these to anticipate 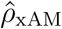, the projected ef-fect correlation estimate purely due to xAM-induced artifact. Across numerous pairs of nominally genetically correlated traits, xAM alone could plausibly produce substantial 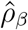 values relative to empirical LDSC estimates, though the relative magnitude of such effects varied considerably across phenotype pairs. Additionally, we provided evidence congruent with a history of xAM at the genetic level by demonstrating that cross-trait even/odd chromosome polygenic score correlations mirror phenotypic cross-mate correlations. Moreover, we showed that cross-trait even/odd chromosome polygenic score correlations explain substantial variation in genetic correlation estimates through their association with cross-mate phenotypic correlations, consistent with the hypothesis that changes to global LD structure induced by previous generations of xAM have resulted in biased genetic correlation estimates.

Taken together, our results show that assortment across phenotypes will bias effect correlation estimates, that cross-mate cross-trait phenotypic correlations among many pairs of phenotypes are strong enough that one or more generations of assortment would substantially inflate effect correlation estimates, and that the joint distribution of causal variants and their estimated effects coincides with what we would expect after one or more generations of xAM. We therefore conclude that xAM provides a previously ignored source of systematic bias in the study of genetic similarity across complex traits, one that precludes the interpretation of the effect correlation estimate as a direct index of biological similarity.

This is perhaps most salient in scientific domains where effect correlation estimates may be interpreted through a clinical lens. In order to better understand the potential consequences of xAM-naïve approaches, we reexamined the results Anttila and colleagues [2], who, in chorus with others [7], interpret substantial effect correlation estimates among psychiatric disorders as evidence of shared pathogenesis that contradicts existing disease classifications. Though our results are agnostic with respect to the adequacy of psychiatric nosology, they illustrate that effect correlation estimates alone do not imply shared pathogenesis and suggest that a substantial fraction of the effect correlation estimates among many pairs of psychiatric traits are largely consistent with xAM alone. Further, modest levels of diagnostic error, when coupled with xAM, can fully account for some previously published estimates (Figures 4c and S11). Reassuringly, our results also demonstrate that particular trait pairs with high rates of comorbidity or characterized by similar symptomatology (i.e., anxiety disorders/major depression and bipolar disorders/Schizophrenia; [26, 37]) exhibit effect correlation estimates greater than what we could plausibly attribute to xAM and diagnostic error. Though it is still possible that such estimates are biased upwards to some extent by either or both factors, the general qualitative interpretation that these disorders share overlapping genetic bases appears robust.

Our results also complicate the interpretation of a number of multivariate analytic frameworks beyond effect correlation estimators. For example, genomic structural equation modeling [6], which takes marker-based genetic correlation estimates as inputs, will propagate xAM induced biases. The consequences for methods such as MTAG [38] are not obvious as xAM will not necessarily decrease prediction *R*^2^ (e.g., see Figure S5b). That is, the apparent gain in predictive performance of multi-phenotype polygenic score aggregates could arise from the global patterns sign-consistent LD induced by xAM. Broadly speaking, our findings mirror recent results regarding the potential impacts of assortative mating across other areas of statistical genetics, including marker-based heritability estimation [17], and Mendelian randomization [19, 39].

There are a number of limitations to the current investigation. Foremost among these are the constellation of assumptions required to predict the expected effect correlation estimate under xAM alone 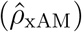 which here reflect a particular model of assortment (see Supplemental Materials S1.1 for further details). Even in the simplified context of two additive traits with uncorrelated effects, anticipating 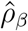 values under xAM alone necessitates numerous unconfirmed assumptions about the dynamics of a particular population, including but not limited to: the panmictic heritabilities, the stability of cross-mate phenotypic correlation structures over succeeding generations, and the stability and transmissability of environmental factors, each of which comprises an active research area [17, 20, 40]. We proceeded under a relatively tractable dynamical framework assuming a fixed number of generations of assortment on two additive phenotypes with constant cross-mate correlations, stable non-heritable sources of variation, and no vertical transmission. Though each of these assumptions is likely to prove untenable for particular trait pairs and thereby compromise the accuracy of our projections, we hypothesize that the general qualitative phenomenon whereby xAM inflates effect correlation estimates is likely to persist for a number of traits. Nonetheless, we caution that these projections are contingent upon a number of consequential decisions, noting that all existing methods are only able to sidestep these decisions by making the strong (and likely incorrect) assumption that mating is random.

Separate from this collection of necessary modeling decisions is the more elemental assumption of *primary phenotypic assortment*–that mate selection is mediated through both the heritable and non-heritable factors underlying a pair of phenotypes. This assumption represents a middle ground between the two extreme scenarios of *genetic homogamy*, in which mates assort exclusively on genotype, and *social homogamy*, where mates assort exclusively on environmental factors [41]. Critically, xAM under pure social homogamy, which is conceptually equivalent to primary phenotypic assortment across 0% heritable phenotypes, would have no impact on either the score correlation or effect correlation estimator (this can be intuited by extrapolating toward 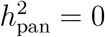 in Figure 2d). However, if pure social homogamy were widespread, we would not expect to observe any association between cross-trait even/odd chromosome polygenic score correlations and cross-mate cross-trait correlations, which was not the case empirically (Figure 5a). To the extent that the assumption of primary phenotypic assortment is violated, it is most likely to be a matter of degree (i.e., heritable and non-heritable phenotype components may be weighted disproporitionately but both impact mate choice to some extent), and can be conceived of as another modeling procedure unlikely to dramatically alter our qualitative conclusions. Still, constructing a generative model that reconciles the association between empirical mating patterns and genetic correlation estimates is a ill-posed inverse problem for which there are multiple solutions, of which we have only explored one.

Finally, though we demonstrate that empirical levels of xAM are substantial enough to have induced systemic bias in the genetic correlation literature, we have not explicitly estimated the extent to which empirical genetic correlation estimates reflect biological versus structural factors. Thus, the development of quantitative methods that jointly model causal loci and their effects (i.e., do not assume *ρ*_*l*_ and *ρ*_*β*_ are equivalent) comprises a prime target for future research. We further remark that xAM is, in essence, a form of population structure, and suggest that the development of such methods may provide insight into residual sources of variation not captured by conventional principal component or mixed-model based correction. Given the increasing evidence that existing methods fail to completely address structural factors, even in ostensibly ancestrally homogenous groups [42, 43], a broader characterization of population structure and new methods for addressing such structure will like be necessary to generate results that are maximally clinically relevant and can be applied equitably.

## 5 Online methods

### 5.1 Simulation framework

We present results derived from two forward-time simulation frameworks: a highly realistic but computationally intensive scheme (hereafter referred to as the *large-scale framework*) and a simplified, computationally efficient scheme (hereafter referred to as the *simplified framework*), both of which we describe in detail in the following section. We justify employing the simplified framework by validating its output against that of the realistic framework, concluding that the simplified framework is sufficient for modeling the effects of xAM. Further, comparing these methods allows us to isolate the impacts (or lack thereof) of several phenomena of interest, including pleiotropy and local linkage disequilibrium (LD). We provide the code necessary to repeat or extend these analyses at https://github.com/rborder/xAM_and_gen_corr. All analyses were conducted using R v4.0.2 or R v4.1.0 [44] unless otherwise stated.

#### 5.1.1 Large-scale framework

With the exception of extending the mating scheme to the bivariate case, we proceded as in [17]. Briefly, we used the genotypes of unrelated European ancestry UK Biobank participants at one million phased, imputed HapMap3 SNPs with minor allele frequency (MAF) ≥ 0.01 as founder population. Meoisis was simulated by dividing the genome into 10 kB blocks and deriving recombination probabilities from a linear interpolation of 1000 Genomes Project Phase 3 recombination map [45], thus preserving the existing local LD structure.

For each simulation, *m*=1e4 loci were selected as causal variants for the two phenotypes *Y, Z* such that their standardized effects *β*_*y*_, *β*_*z*_ were jointly bivariate Gaussian with respective variances 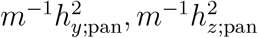 and correlation *ρ*_*β*_. Phenotypes were then generated according to the additive model in (1). Individuals (represented by pairs of phenotype values) were mated according to the exchangeable cross-mate correlation regime described in Section 5.1.3.

In order to avoid any potential confounds due to within-sample relatedness, each coupling produced a single offspring, thereby halving the sample size with each successive generation and ensuring no pairs of individuals share any indentical-by-descent segments with respect to the founder population. At each generation, we obtained heritability and genetic correlation estimates from HE regression as implemented in GCTA v1.93.2b [28], LDSC regression as implemented in LDSC v1.0.1 [3], and REML as implemented in BOLT-LMM v2.3.4 [29]. For LDSC, we estimated LD scores within sample using input paramater `ld-wind-cm=1.0` and obtained GWAS summary statistics using plink2 v2.0 [46]. To improve computational efficiency, we analyzed subsamples of *n*=4e5 individuals, with the exception of the fourth and fifth generation results presented in Figure 2a, which were respectively limited to *n*=2e5, 1e5 by the diminishing sample size at each generation.

#### 5.1.2 Simplified framework

Given the computational resources required by the large-scale framework in light of the large number of trait pairs we wanted to base simulations on (each run of the large-scale framework generates several terrabytes of output), we developed a simplified framework using entirely synthetic data and only including causal loci. We constructed founder genotypes by randomly drawing 2*m* haploid SNPs with *m* allele frequencies distributed uniformly on [0.0, 0.99] independently for *n* individuals, with meiosis proceeding as described in Section 5.1.2. Any potential confounds due to relatedness were avoided as in the large-scale framework described above.

##### Phenotype simulation with and without pleiotropy

We again generated phenotype pairs according to the additive model (1), restricting our focus to the case of genetically orthogonal traits. Given a specified effect correlation (in this case *ρ*_*β*_ = 0), pleiotropy is irrelevant. We demonstrated this by simulating phenotypes under either complete pleiotropy (all causal variants have non-zero effects on both phenotypes) and zero pleiotropy (all causal variants have non-zero effects on only one of pair of phenotypes).We independently sampled genetic effects *β*_*y*_, *β*_*z*_ from the univariate Gaussian distributions with respective variances 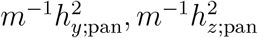, using a single set of *m* causal variants to simulate complete pleiotropy and two independent sets of *m*/2 causal variants for to simulate zero pleiotropy. Figure S4 demonstrates that, as expected, the fraction of causal variants shared between phenotypes has no bearing on the impact of xAM. However, as the zero pleiotropy simulation procedure reduced sampling variance across simulations (in a finite sample, *ρ*_*β*_ has greater variance when causal variants overlap), we utilized this approach for the results presented in Section 2.4. The number of causal variants had no apparent impact on quantities of interest (Figure S1).

##### Manipulating local LD

Denoting the first and second copies of an individual’s haploid genotypes at the jth diploid locus by 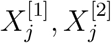, respectively, we manipulated local LD during meiosis I by enforcing recombination events at each single line with probability *p*_recomb_. and each double line with probability 0.5 below:

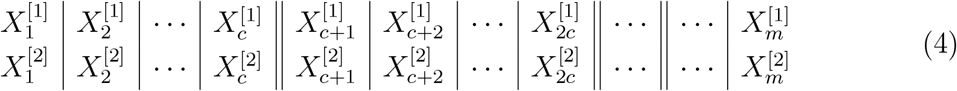

Here *c* = *m*/20 divides the genome into 20 independently inherited “chromosomes”. Within each chromosome, recombination events occurs between contiguous loci with probability *p*_recomb._ ϵ (0.0, 0.5], such that *p*_recomb._=0.5 corresponds to unlinked loci, with the strength of local LD in the subsequent generation increasing as *p*_recomb._ → 0. Figure S3 demonstrates the irrelevance of *p*_recomb._ with respect to liability correlation and estimated effect correlation; simulations with strong local LD (*p*_recomb._ = 0.01) and weak local LD (*p*_recomb._ = 0.5) yielded results consistent with the realistic framework simulation results.

#### 5.1.3 Mating regimes

For two phenotypes *Y, Z*, denote their joint distribution across mates by

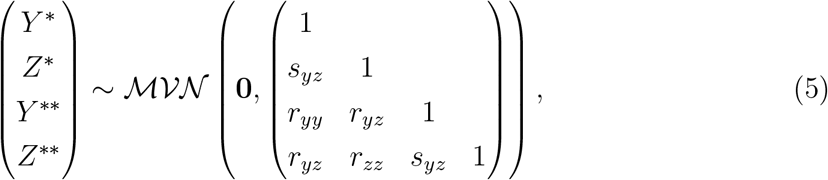

where the number of asterisks distinguish the two mates. Whereas the within-mate single-trait correlation (i.e., the conventional phenotype correlation) *s*_*yz*_ is unaffected by mating patterns within a given generation, the cross-mate single-trait correlations *r*_*yy*_, *r*_*zz*_ and the cross-mate cross-trait correlation *r*_*yz*_ are free parameters determined by the mating regime.

For the large-scale simulations presented in Section 2.3, individuals were mated by randomly splitting the sample in half and pairing individuals across the two subsamples, ordering each subsample on a linear combination of their phenotypic values and Gaussian noise. This corresponds to an exchangeable cross-mate correlation structure with all cross-mate correlations *r*_*yy*_, *r*_*yz*_, *r*_*zz*_ equal to a single input parameter *r*_mate_. Though this method is computationally efficient and thus well-suited to this particular use-case, it is incapable of achieving arbitrary cross-mate correlation structures. Thus, a more flexible approach was required for the simulations presented in Section 2.4, which were based on empirical estimates of the cross-mate correlation parameters.

We achieved arbitrary cross-mate correlation structures, a problem for which we were unable to find a previously published solution, by using an ad hoc approach based on propensity-score matching methods as implemented in the R package *MatchIt* v4.2.0 [47]. Specifically, we randomly split the sample in half and used the nearest Mahalanobis distance matching algorithm on a scalar multiple of each of the two phenotypes, their product, and their difference. The multiplier hyperparameter was chosen numerically by finding the rational function of the correlation parameters in (5) that minimized the average *l*_∞_ distance between the desired and achieve cross-mate correlations across replicate samples. Discrepancies between target correlations and achieved correlations were modest: across all generations of UKB simulations, the average median discrepancy was 0.0007 and 90% of all cross-mate correlations were within 0.02 of their target values (see Tables S3 and S5). We further validated this approach by directly comparing it to the linear combination ordering approach described above, specifying a single value for each of the parameters *r*_*yy*_ = *r*_*zz*_ = *r*_*yz*_ = *r*_mate_. Both methods yielded equivalent results (Figure S2).

### 5.2 Phenotyping

We examined a broad array of phenotypes across two large cohorts for which cross-mate correlation estimation was feasible. This included 341,997 unrelated European ancestry UK Biobank participants [21] and 373,283 spousal pairs drawn from Danish registry data [22–25].

We *a priori* selected a variety of previously-studied phenotypes discussed in two influential publications [2, 3], discarding measures redundant with available measures of higher quality. For example, in the UK Biobank, we excluded college completion (`f.6138`) which factors into the higher-resolution measure of years of education, while retaining the latter. Similarly, we excluded total cholesterol (`f.30690`) while retaining the component phenotypes LDL cholesterol (`f.23405`), HDL cholesterol (`f.23405`), and triglycerides (`f.30870`). Blood biochemistry phenotypes were further adjusted for statin usage.

### 5.3 Identification of mating pairs in the UK Biobank

We identified putative mate pairs in the UK Biobank using a procedure broadly similar to that of Howe and colleagues [48] with two distinctions: 1. we incorporated geographical information to impute cohabitation status, and 2. we relaxed the condition that both potential mates must report having lived in the same location for precisely the same number of years. We considered sex-discordant pairs of European ancestry participants from the same assessment centers (`f.54`) who reported living with a spouse (`f.699`), excluding pairs discordant on any of the following measures:

- latitude / longitude of home location rounded to the nearest kilometer (`f.20074, f.20075`)
- inverse distance between home and nearest road / major road (`f.24010, f.24012`)
- coastal proximity (`f.24508`)
- household size (`f.709`)
- number of vehicles (`f.728`)
- accommodation type (`f.670`)
- rental status (`f.680`)

To minimize the possibility of identifying cohabitating relatives, we required pairs to be discordant on the age of at least one parent (`f.1807, f.1846, f.2946, f.3526`) and removed third degree or closer relatives as via estimated kinship coefficients (see Supplemental Materials S1.3.1 for kinship estimation details). Finally, we removed all participant groupings meeting these criteria including more than two individuals. This resulted in a total of 40,697 putative mate pairs (i.e., 81,394 individuals).

### 5.4 Identification of mating pairs in the Danish population cohort

We obtained empirical estimates of spousal correlations for five psychiatric disorders in the Danish Population using the Danish Civil Registration System [23, 24], the Danish National Patient Register [25], and the Danish Psychiatric Central Research Register [22]. We first randomly selected 500,000 individuals born between 1981 and 2005 from the Danish Civil Registration System. The parents of these individuals served as our sample of mates (373,283 spousal pairs total).

### 5.5 Cross-mate phenotypic correlation estimation

We estimated cross-mate correlations using a structural equation modeling approach via the R package *lavaan* v0.6-8 [49]. Specifically, for the UK Biobank phenotypes, we estimated the sex-constrained structural model corresponding to Eq. (5), with *Y* and *Z* representing a pair of phenotypes after regressing out the effects of sex, age, sex×age and age^2^ (with the exception of metabolic phenotypes, which were also adjusted for statin use). For the psychiatric phenotypes, which were dichotomous, we proceded similarly, using the corresponding liability-threshold models to estimate cross-mate tetrachoric correlations. All models were estimated via maximum likelihood with asymptotic standard errors and using pairwise complete observations.

### 5.6 Estimation of GWAS summary statistics in the UK Biobank

For each phenotype of interest, we estimated association regression weights at 1,157,133 imputed HapMap3 SNPs with missingness < 0.01, Hardy-Weinberg equilibrium *p* > 1e-5, INFO imputation quality score > 0.9, and MAF > 0.01 using plink v2.0 [46]. Covariates included sex, age, sex×age, age^2^, 21 genomic principal components, assessment center, and genotyping batch (though we note that metabolic phenotypes were further adjusted for statin use as detailed above). Analyses were restricted to a subset of 341,997 unrelated European ancestry individuals (see Supplemental Materials S1.3.1 for details of relatedness estimation).

### 5.7 Empirical heritability and genetic correlation estimates

#### 5.7.1 UK Biobank cohort

We estimated marker-based heritabilities and genetic correlations using LDSC v1.0.1 [30] with internal summary statistics (see previous section) and LD scores (computed for regression SNPs with input parameter `ld-wind-cm 1.0`). We obtained pedigree-based heritability estimates from the empirical literature, preferring estimates derived in UK-based samples whenever available. In the case of classical twin studies, we used results from “ACE” models, which are expected to yield downward-biased heritability estimates under AM [35] and thus constitute the more conservative option, when available (Table S2).

#### 5.7.2 Literature-derived estimates for psychiatric disorders

We used a subset of the LDSC heritability and effect correlation estimates reported by Anttila and colleagues [2] corresponding to the set of psychiatric disorders for which we estimated cross-mate cross-trait correlations in the Danish population cohort. As with the UK Biobank phenotypes, we extracted pedigree-based heritability estimates from demographically comparable samples preferring “ACE” estimates when available (Table S4).

### 5.8 Projected genetic correlation estimates under xAM alone

We implemented a simulation based approach to quantify the magnitudes of empirical 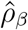 estimates with respect to expectations for genetically orthogonal traits under xAM alone. For each pair of traits *Y, Z* we ran 400 replicate simulations with a founder population size of *n*=64,000, setting the panmictic heritability parameters to empirical marker or pedigree-based heritability estimates and allocating *m* = 1000 distinct causal variants for each phenotype (thus ensuring *ρ*_*β*_ = 0). We then simulated three generations of xAM using empirical estimates of the cross-mate correlations 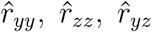, at each generation obtaining method-of-moment estimates of *ρ*_*β*_, which we denote 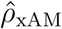, using HE regression, which, as demonstrated in Figures 2a to 2d, produces results equivalent to LDSC regression.

In order to propagate uncertainty in the empirical estimates through the simulations, simulation inputs were randomly sampled from the asymptotic sampling distributions of the corresponding empirical estimates. We first randomly sampled the empirical genetic correlation estimate 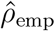 and the input parameters 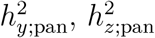, *r*_*yy*_, *r*_*zz*_, *r*_*yz*_ from their joint sampling distribution

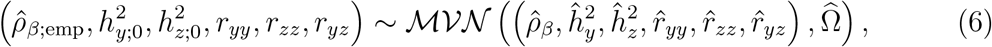

where 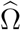 denotes the empirical variance-covariance matrix. We then compared the 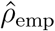 to the estimate projected by the simulation, which we denote 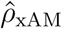, to compute the ratio statistic 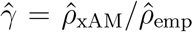. The empirical quantiles of 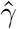 which approximate its posterior distribution, were then used to compute credible intervals. For each trait pair, we produced two sets of results, once substituting pedigree-based heritability estimates 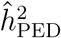 for the panmictic heritability inputs and again substituting marker-based heritability estimates 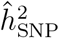. Unfortunately, the sample variance-covariances of 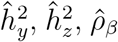 are not reported by the LDSC software and were not available for any of the literature-derived estimates, and were treated as independent by necessity. That is, only the off-diagonal elements of 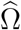 corresponding to the cross-mate correlation parameters, which we estimated via structural equation modeling, were nonzero.

### 5.9 Cross-chromosome correlation of polygenic scores in the UK Biobank

We first randomly split the UK Biobank into two disjoint subsamples, with 8 0% of participants comprising the training set and the remaining 20% comprising the test data. Next, we ran a GWAS for each phenotype in the training sample using covariates and procedures identical to those described in Section 5.6 and clumped results using plink v1.90b6.21 using a 250kB sliding window and an *R*^2^ threshold of 0.05. We then used the training sample summary statistics to compute polygenic scores for separately for even and odd chromosomes using up to 20 *p*-value thresholds spaced logarithmically on the interval [5e-8, 1.0], with the resulting number of scores depending on the maximally significant SNP f or a given trait.

For any given pair of phenotypes *Y, Z*, there are two potential even/odd chromosome polygenic score correlations, 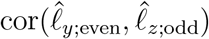 and 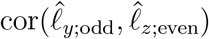, neither of which is of greater interest than the other. Thus, we estimated both quantities via a single parameter, *ρ*_*l*;eo_, in the constrained structural model

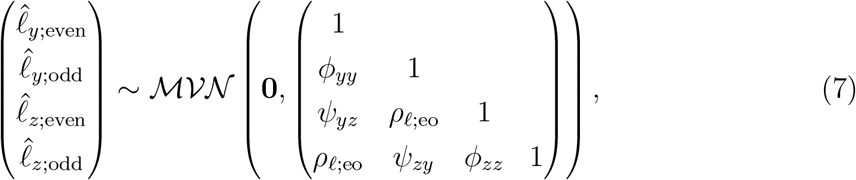

where *ρ*_*l*;eo_ is the only free parameter of interest. For the simpler case of single-trait even/odd correlations, we used the correspondingly simple bivariate Gaussian structural model. Again, we estimated structural models using the R package *lavaan* v0.6-8 [49] via maximum likelihood with asymptotic standard errors.

Finally, we evaluated the relationships between even/odd chromosome polygenic score correlation estimates and both cross-mate cross-trait correlation estimates and genetic correlation estimates across all pairs of UKB sample phenotypes. We examined these associations in the context of naïve linear models, which don’t account for heteroskedasticity and sampling error in the predictor, but also using a Bayesian measurement error model as implemented in the R package *brms* v1.8 [50], which does.

## S1 Supplemental material

### S1.1 Theory of assortative mating

#### S1.1.1 Long range sign-consistent LD

Denote quantities relating to the members of a parent-parent-offspring trio by [·]*, [·]**, 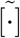, respectively. Let 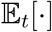 denote the expectation of a quantity after *t* generations of positive xAM. Let *X*_*i*_, *X*_*j*_ denote mean-deviated trait-increasing-allele counts at two causal loci for mean-deviated phenotypes *Y, Z*, respectively. We assume that *X*_*i*_, *X*_*j*_ are unlinked at panmixis (when *t* = 0). For simplicity, we assume each variant is causal for one and only one of the two phenotypes, i.e.,

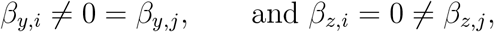

and we assume the cross-mate covariances are positive and symmetric such that for all *t* ≥ 0

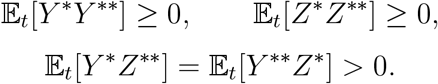

We assume *primary phenotypic assortment*: mates’ genotypes are conditionally independent given the heritable component of either mate’s phenotype. Denoting the heritable components of *Y, Z* by *l*_*y*_, *l*_*z*_, respectively, we assume

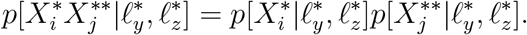

Denote the conditional expectation of an individual’s causal variant genotype at locus *i* given the heritable components of their phenotypes by

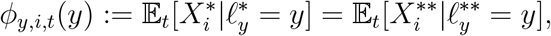

and denote the conditional expectation of an individual’s causal variant genotype given their mate’s heritable phenotype components by

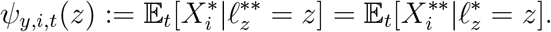

We assume that *ϕ*_*i,t*_(*y*),*ψ*_*i,t*_(*z*) are monotone functions in their respective arguments that cross the origin and that agree in sign with the effect of locus *i* such that

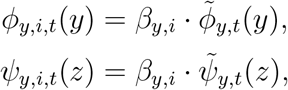

where 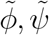 are non-trivial monotone increasing functions that cross the origin. That is, we assume that for a *trait-increasing* allele for *Y* (i.e., when *β*_*y,i*_ > 0) *ϕ*_*y,i,t*_(*y*) is an *increasing* function, and further, as *Y* and *Z* are positively correlated across mates, *ψ*_*y,i,t*_(*z*) will also be increasing. Likewise, if *X*_*i*_ is *trait-decreasing ϕ*_*y,i,t*_(*y*) and *ψ*_*y,i,t*_(*z*) will be *decreasing* in their respective arguments. Though rather technical in specification, these assumptions are intuitive: if I know one mate is “high” with respect to (the heritable component of) phenotype *Y*, I expect “higher” genotypic values at their own loci and their partner’s loci that increase *Y* and “lower” genotypic values at their own and their partner’s loci that decrease *Y*. Further, this is a weaker assumption than those of previous authors (e.g. in the single-trait case [51] assumes linearity such that *ϕ*_*y,i,t*_(*y*) ∝ *β*_*y,i*_ · *y*).

Having introduced our assumptions, we seek to investigate the linkage disequilibrium between *X*_*i*_, an arbitrary causal variant for phenotype *Y* and *X*_*j*_, an arbitrary causal variant for phenotype *Z*, after *t* > 0 generations of positive xAM across *Y* and *Z*. At every generation, we can factor the covariance 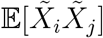 using the possible patterns of inheritance:

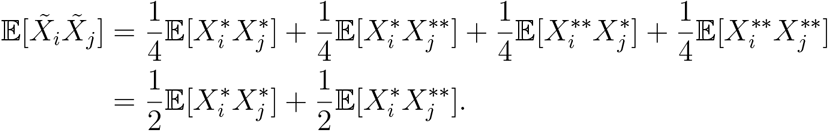

As by definition

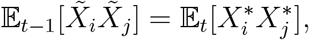

this induces the following recurrence relation:

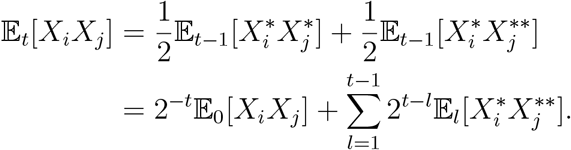

By hypothesis, *X*_*i*_, *X*_*j*_ are unlinked at panmixis and thus 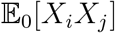 is zero. That leaves us with the sequence of cross-mate cross-locus moments 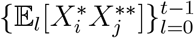 terms.

Applying the assumption of primary phenotypic assortment, we have

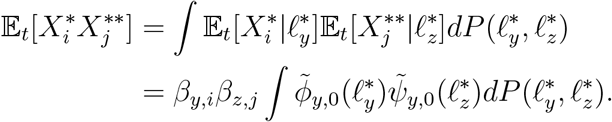

The above integral is strictly positive, thereby yielding

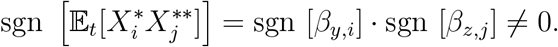

All together, we have

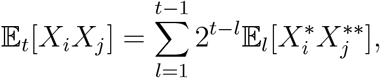

where each of the terms in the above sum has sign sgn [*β*_*y,i*_] sgn [*β*_*z,j*_], establishing that

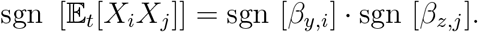

#### S1.1.2 Inflation of GWAS statistics

We now show that assortment on a single trait (sAM) leads to inflated GWAS effect estimates. This is a simplification of the bivariate case.

By the same argument used in the previous section, it is easy to see that sAM on phenotype *Y* induces sign-consistent long-range LD such that

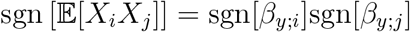

for all causal variants indexed *i, j*. Without loss of generality, assume all variants *i, j* ϵ {1, … , *m*} are causal and denote the correlation between standardized variants at loci index *i, j* by

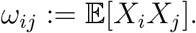

We estimate the GWAS effect of variant *X*_*i*_ on *Y* as

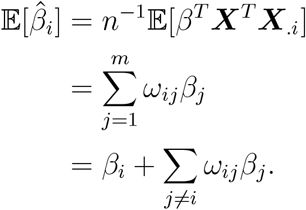

Each of the above summands has sign

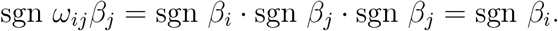

Thus, when *β*_*i*_ > 0 we have 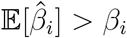 and when *β*_*i*_ < 0 we have 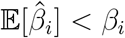. That is, the magnitude of the GWAS effect estimate is inflated upwards.

#### S1.1.3 Correlation of even/odd chromosome cross-trait polygenic scores

We now consider the correlation between polygenic scores for *Y* and *Z* constructed on disjoint sets of loci (as in even/odd chromosome polygenic score correlations). Partition the indices of the genome into two non-empty sets 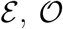, such that {1, … , *m*} is their disjoint union. Define the estimated polygenic scores

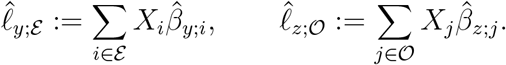

These correspond to a polygenic score for *Y* built only on even chromsomes and a polygenic score for *Z* built only on odd chromsomes. The expected cross-trait even/odd correlation polygenic score covariance is then equal to

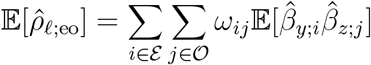

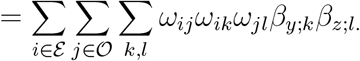

Compare this to the true cross-trait even/odd chromsome polygenic score correlation

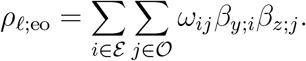

Pulling this out of the previous expression yields

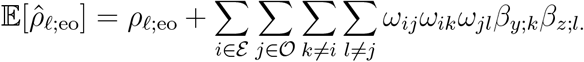

Dealing with the above summand requires fulling understanding the relationship *β*_*y;i*_, *β*_*z*;*j*_ and *ω*_*ij*_ for all *i, j*, which is beyond the scope of the current manuscript. For now, we address the simpler case of the correlation between polygenic scores for a single phenotype *Y* restricted to disjoint sets of chromosomes as analyzed in [18]. Again, we assume that 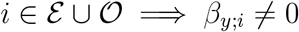 for simplicity. In this case, we have

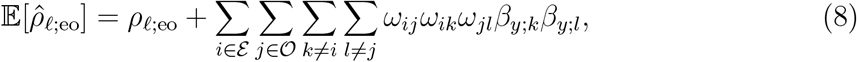

where each of the above summands has sign

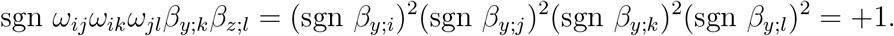

Likewise, the summands comprising the true correlation

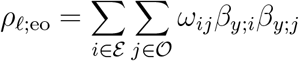

also all have sign +1, altogether yielding the chain of strict inequalities

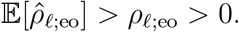

That is, as anticipated by Yengo et al. [18], sAM induces a true correlation between genetic liabilities restricted to disjoint collections of loci. Note that we do not require the assumption of equilibrium.

### S1.2 Supplementary simulation study results

#### S1.2.1 GWAS effect estimates are biased upwards under xAM

We performed additional simulation studies examining the impact of xAM on GWAS effect estimates by comparing estimated regression slopes to their true values at each generation in the context of the simplified forward-time simulation framework described in the Online Methods. In contrast to effect correlation estimators, on which single-trait AM (sAM) has limited effects [5, 52], association statistics are biased upwards in magnitude under sAM as each causal variant weakly will tag every other causal variant (see S1.1.1 and S1.1.2). We therefore sought to distinguish the effects of sAM and xAM by varying the cross-mate cross-trait correlation *r*_*yz*_ independently of the crossmate single-trait correlations *r*_*yy*_ and *r*_*zz*_. Specifically, we consider two phenotypes *Y*, *Z*, each with *m* = 2,000 non-overlapping causal variants and panmictic heritabilities equal to 0.5. Letting asterisks distinguish mates, we varied the cross-mate correlation parameters governing assortment

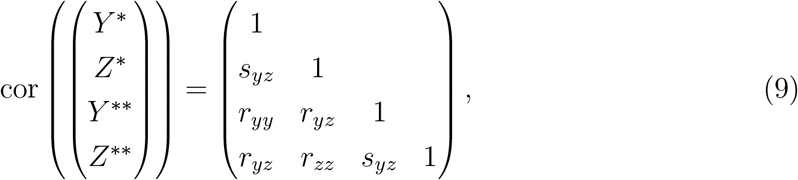

fixing *r*_*zz*_ ≡ 0.5 and varying (*r*_*yy*_, *r*_*yz*_) ϵ {0.0, 0.25, 0.5}^2^, running 100 simulation replicates per experimental condition. Focusing on *Y*, we have sAM when *r*_*yy*_ > 0 and *r*_*yz*_ = 0 and xAM when *r*_*yz*_ > 0

At each generation, we emulate a GWAS by regressing *Y* on each of the 2*m* standardized genotypes to estimate *β*_*j*_, *j* = 1, … , 2*m* in a sample of size *n* =32,000. We then report two metrics: the average magnitude of GWAS slope error relative to the average effect size 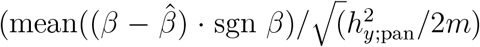; hereafter termed *slope inflation*), which we present in Figure S5a, and the correlation between the true liability *l*_*y*_ = Σ_*i*_ *X*_*i*_ *β*_*i*_ and estimated polygenic score 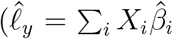; hereafter termed *score correlation*), which we present in Figure S5b.

As expected, we observed that the strength of single-trait correlation biases GWAS effects upwards in magnitude regardless of the cross-trait correlation (e.g., after three generations, the average slope inflation per 0.05 increase in *r*_*yy*_ was 0.143 [*se*=0.0374%]). We also saw that cross-mate cross-trait correlation alone causes upwards inflation and exacerbates the inflation caused by single trait correlations (e.g., after three generations, the average slope inflation per 0.05 increases in *r*_*yz*_ were 0.157% [SE 0.0097%] and 0.143% [SE 0.0310%] for *r*_*yz*_=0 and *r*_*yz*_>0, respectively). To summarize, both assortment on a single trait and assortment across multiple traits upwardly bias GWAS effect estimates in subsequent generations, and these biases are exacerbated when both forms of assortment are present.

Polygenic score results are less straightforward as single-versus cross-trait assortment exert countervailing influences on the correlation between the true and estimated polygenic scores. Regressing the difference between the score correlations for phenotype *Y* after three generations of assortment and those at generation zero (panmixis), the difference score increases by 3.46e-3 (SE: 2.59e-5) per 0.05 increase in *r*_*yy*_ and decreases by 1.10e-3 (SE: 2.59e-5) per 0.05 increase in *r*_*yz*_. We interpret this to mean that single-trait assortment, even though it increases the average effect estimate error, leads to a stronger correlation between the true and estimated polygenic scores, presumably because all causal variants tag one another. On the other hand, cross-trait assortment, which induces correlations between causal variants for one trait and causal variants for another, inflates GWAS slope estimates while decreasing the score correlation, at least when the two traits are genetically orthogonal.

#### S1.2.2 xAM exacerbates the impacts of misclassification errors on effect correlation estimates

We ran an additional set of simulations again using the simplified forward-time simulation framework to characterize the impact of xAM on binary traits subject to misclassification. We again simulated two continuous phenotypes with no pleiotropy, this time fixing all the cross-mate correlation parameters to 0.5 while varying trait prevalence as well as the rate and directionality of misclassification errors. Prior to effect correlation estimation, we dichotomized phenotypes using the cutoffs derived from the standard normal quantile function. We introduced misclassification errors by randomly selecting a fixed number of true cases for the first phenotype and relabeling them as controls for the first and cases for the second, an preceding analogous for the second phenotype. As anticipated, misclassification errors induced artifactual 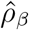 estimates among genetically orthogonal traits and this inflation was in turn exacerbated by the extent of xAM across the two traits (Figure S6). Further, in contrast to xAM, the impacts of misclassification errors depended on trait prevalence such that classification-induced 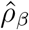 inflation was more pronounced for less common binary traits (Figure S6).

### S1.3 Psychiatric phenotype definitions

The Danish Civil Registration System has been registering all people legally residing in Denmark since 1968, and it includes information about sex, date of birth, parental links, and life events (e.g., migration or death). The System is linked via anonymized identification numbers to the Danish National Patient Register and the Danish Psychiatric Central Research Register that include all diagnostic information regarding general medical conditions and specific psychiatric conditions, respectively, including all inpatient and outpatient contacts. We estimated estimated cross-mate tetrachoric correlations within and across five psychiatric phenotypes considering both ICD-8 and ICD-10 definitions (Table S1). The definitions are based on those by the iPSYCH initiative [53].

**Table S1:** Psychiatric phenotype definitions

|  | Phenotype | ICD-8 code | ICD-10 code |
| --- | --- | --- | --- |
| **ANX** | Anxiety disorders | NA | F40.0-F40.2,<br>F41.0-F41.1, F42,<br>F43.0-F43.1 |
| **ADHD** | Attention deficit/<br>hyperactivity disorder | NA | F90.0 |
| **BIP** | Bipolar disorders | 296.19, 296.39, 298.19 | F30-F31 |
| **MDD** | Single and recurrent<br>depressive disorder | 296.09, 296.29,<br>298.09, 300.49 | F32-F33 |
| **SCZ** | Schizophrenia | 295.x9 (excl 295.79) | F20 |

#### S1.3.1 Identification of unrelated European-ancestry individuals in the UK Biobank

From among 488,363 UK Biobank participants, we retained putative “White British” individuals using field f.22006 (*n*=409,692). We then filtered out 199 individuals with excess genotype missingness (>0.05), 312 individuals with a mismatch between self-reported and genetic sex, 999 inviduals with excess heterozygosity (≥5 standard deviations above the mean), and 90 individuals who requested their data be redacted. We then removed 629 individuals related to ten or more individuals (KING coefficient ≥ 2^−9/2^) as a preprocessing step to the application of the `maximal_independent_set` algorithm implemented in the NetworkX Python package [54]. This resulted in 342,257 unrelated individuals. In contrast to [21], who estimated kinships using ≈92,000 common SNPs with small loadings onto the first few PCs in the full sample (including multiple ancestries; see S3.7 of [21]), we esitmated kinships using 561,780 common SNPS in a sample of European ancestry individuals. The close relatives the UKB identified in `f.22021` are a subset of our more conservative approach: we identified all 81,218 related individuals in this subsample identified by the UKB plus an additional 3,261 not identitified by [21].

## Supplementary Figures

**Figure S1:**
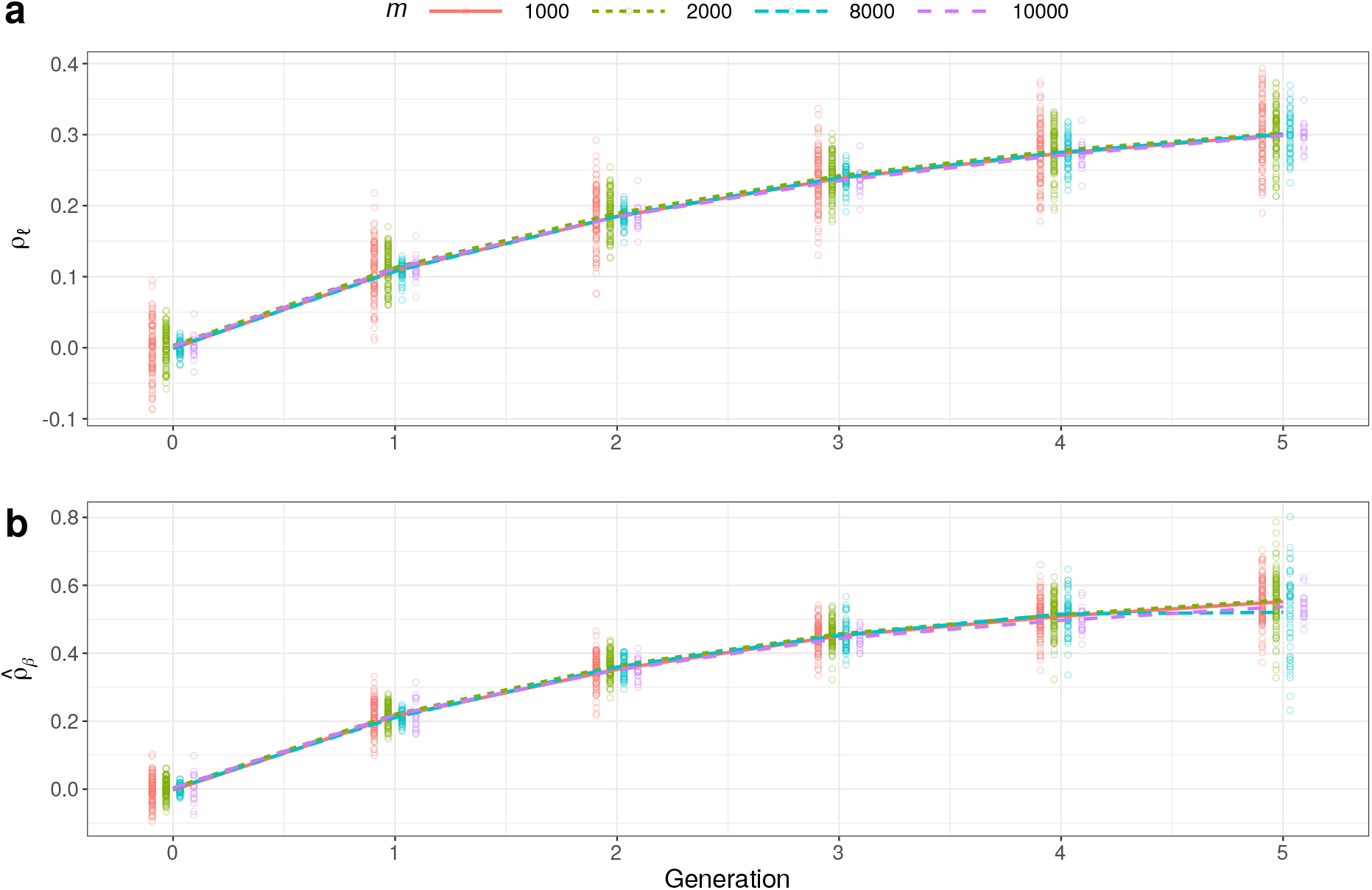
Varying the number of causal variants has no impact on (a) true score correlations or (b) estimated effect correlations across simulations of genetically orthogonal traits with panmictic heritabilities and cross-mate correlations fixed at 0.5.

**Figure S2:**
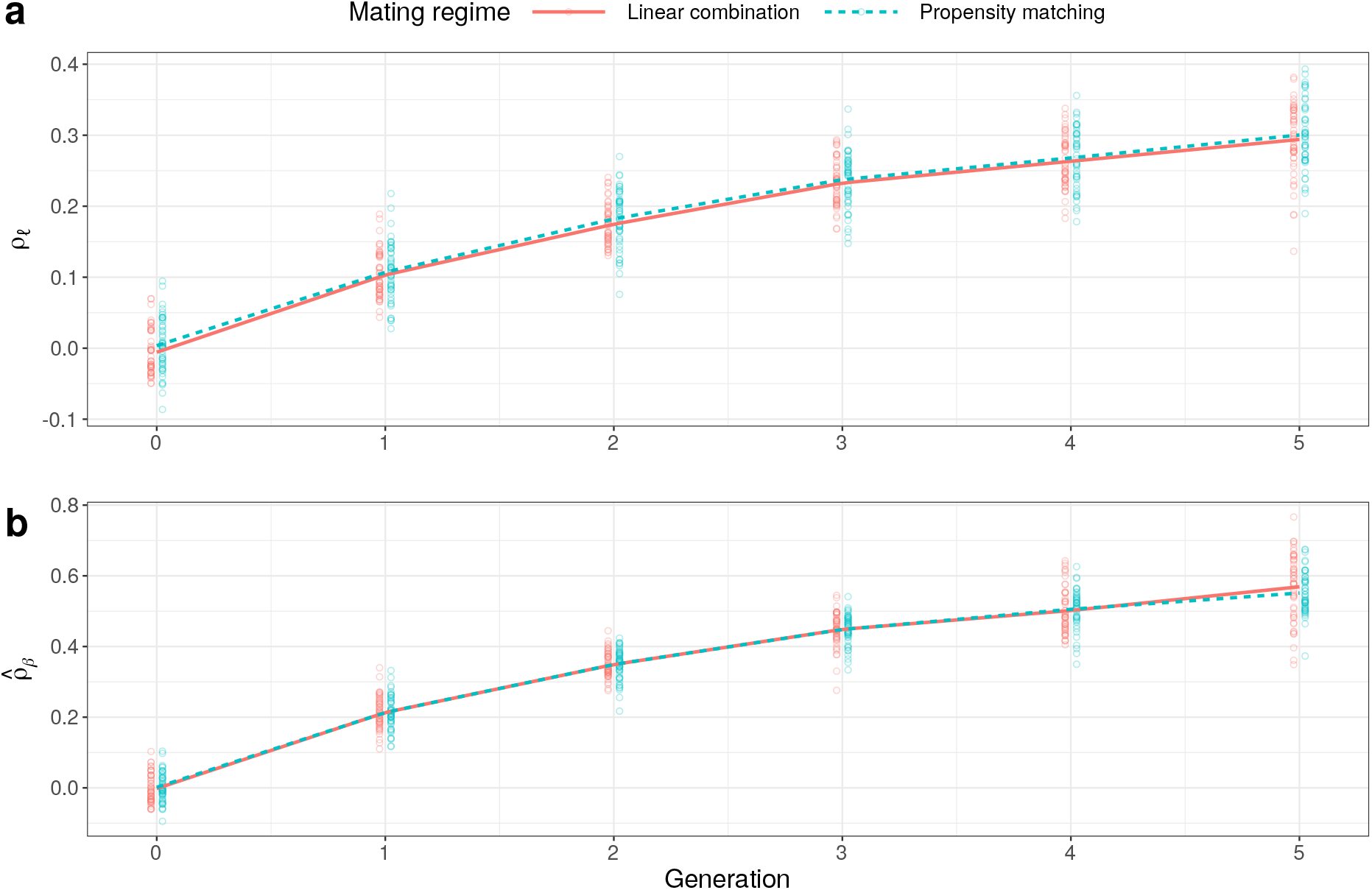
Equivalence of mating regimes under exchangeable correlation structure for genetically orthogonal traits with respect to (a) true score correlation or (b) estimated effect correlation. Panmictic heritabilities and cross-mate correlations were fixed at 0.5 across simulations.

**Figure S3:**
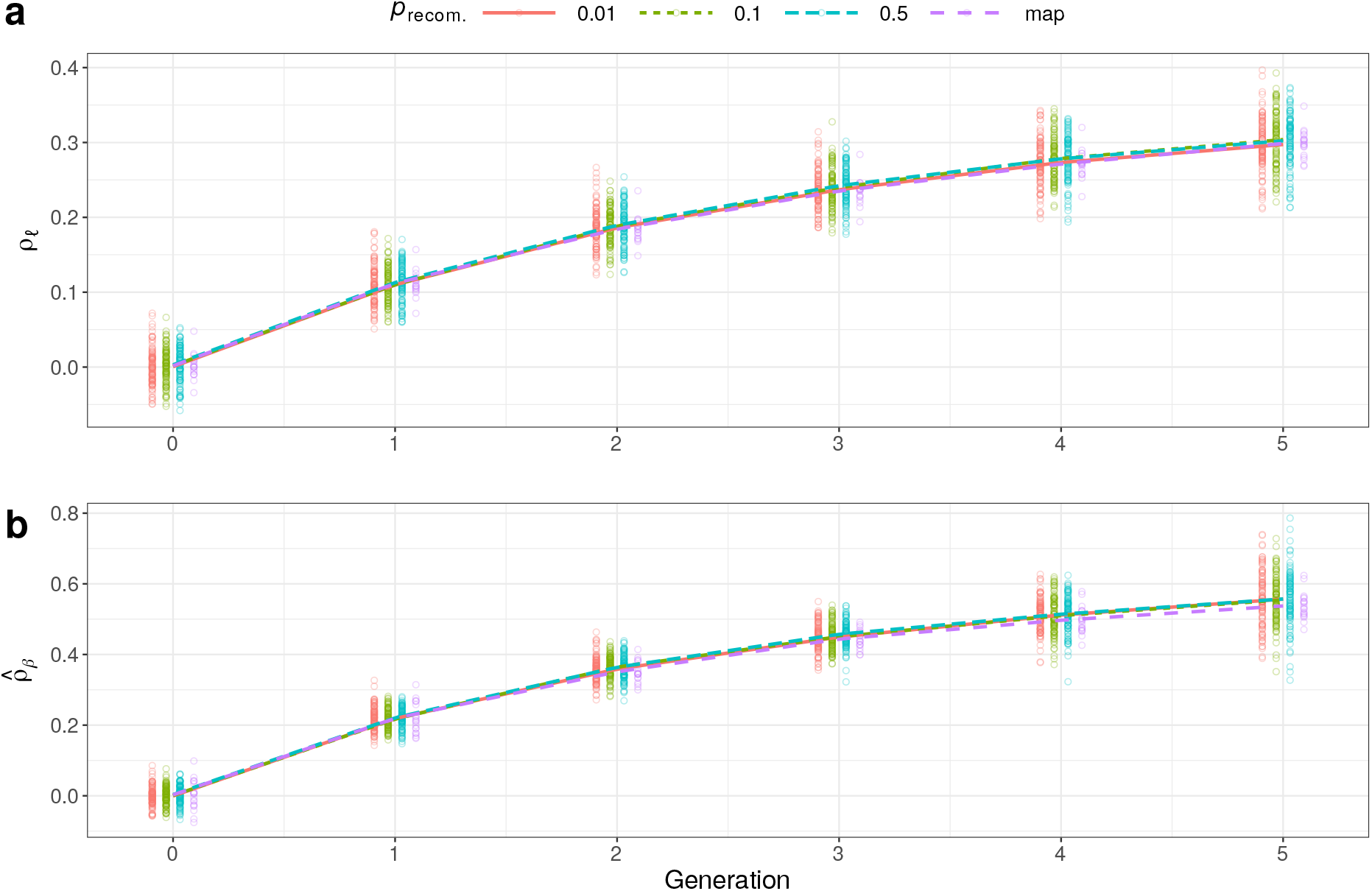
Impact of local LD in synthetic data for genetically orthogonal traits with panmictic heritabilities and cross-mate correlations fixed at 0.5. Enforcing recombination probabilities between continguous loci (*ρ*_recom._) at varying fixed values or using an empirical recombination map has no impact on the (a) true score correlation or (b) estimated effect correlation across simulations.

**Figure S4:**
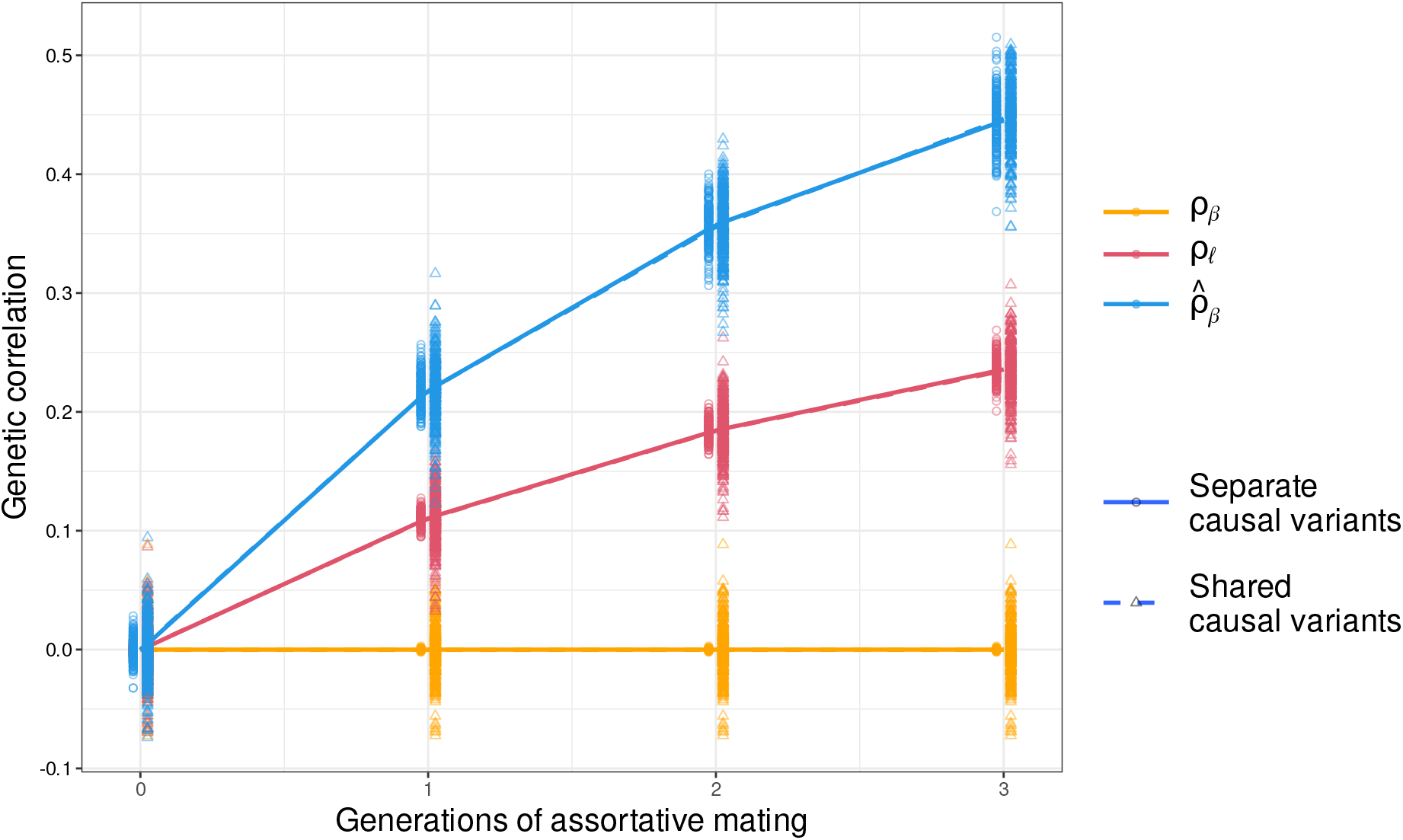
Score correlation and estimated effect correlation for genetically orthogonal traits subject to xAM with and without pleiotropy under orthogonal effects. Across simulations, panmictic heritabilities and cross-mate were correlations fixed at 0.5. Under the separate casual variants regime, *β*_*y*;*i*_ ≠ 0 if and only if *β*_*z*;*i*_ = 0 for each causal variant indexed *i*, whereas under the shared causal variants regime, every variant is causal for both *Y* and *Z* but the effects are drawn independently.

**Figure S5:**
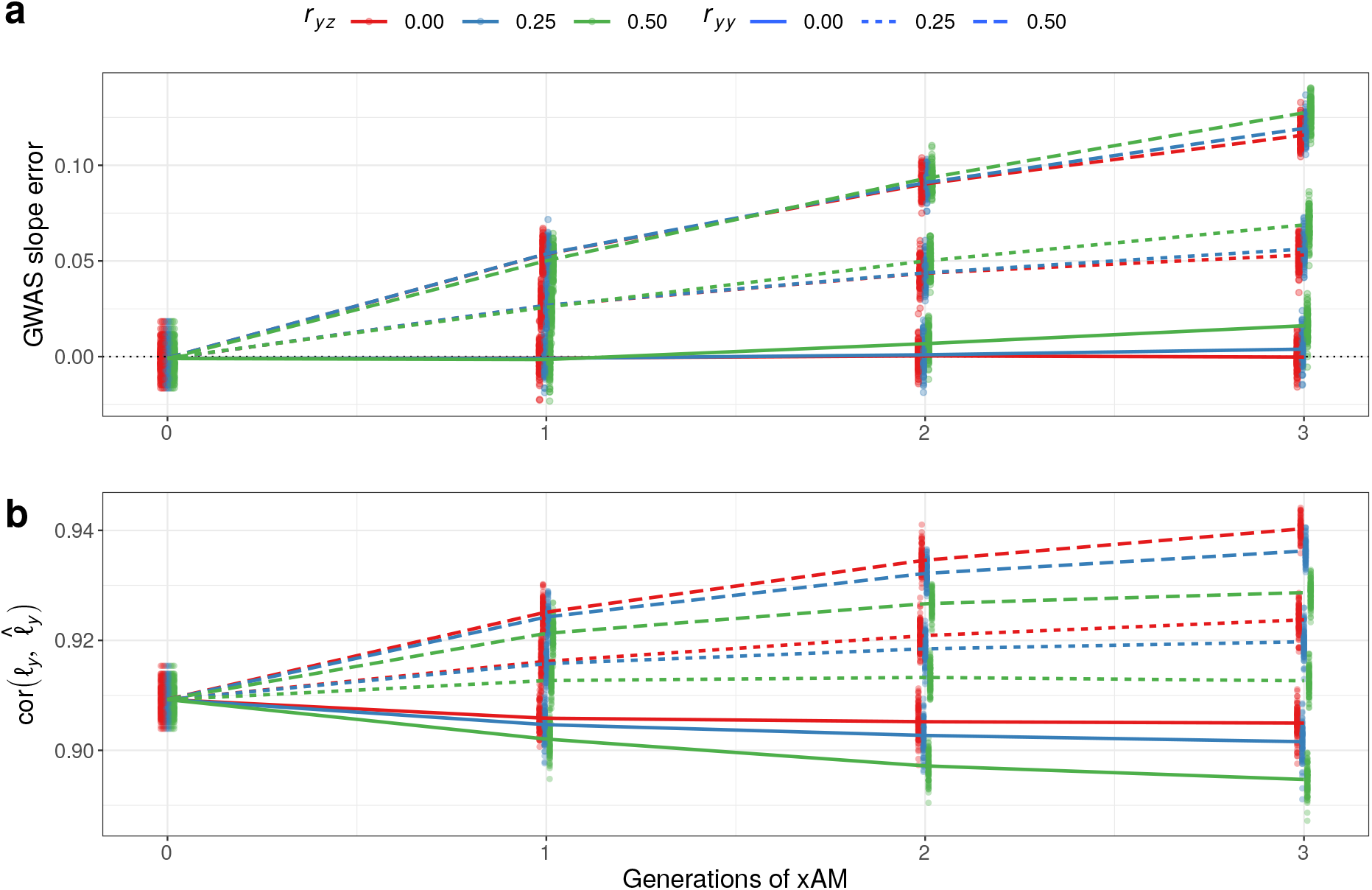
Impact of assortative mating on GWAS regression weights and polygenic scores. Across all simulations, the panmictic heritabilities for *Y* and *Z* are set at 0.5, the true effect correlation is fixed at 0.0, and the cross-mate correlation with respect to *Z*, *r*_*zz*_, is fixed at 0.5, whereas *r*_*yy*_ and *r*_*yz*_ vary across experimental conditions. (a) Slope inflation of GWAS estimates with respect to *Y* under positive xAM depends strongly on *r*_*yy*_ and modestly on *r*_*yz*_. (b) Correlation between the true and estimated polygenic scores for *Y* is inflated by *r*_*yy*_ and deflated by *r*_*yz*_. Intuitively, increasing *r*_*yy*_ induces sign-consistent LD among causal variants for *Y* whereas increasing *r*_*yz*_ induces sign-consistent LD across causal variants for *Y* and *Z*

**Figure S6:**
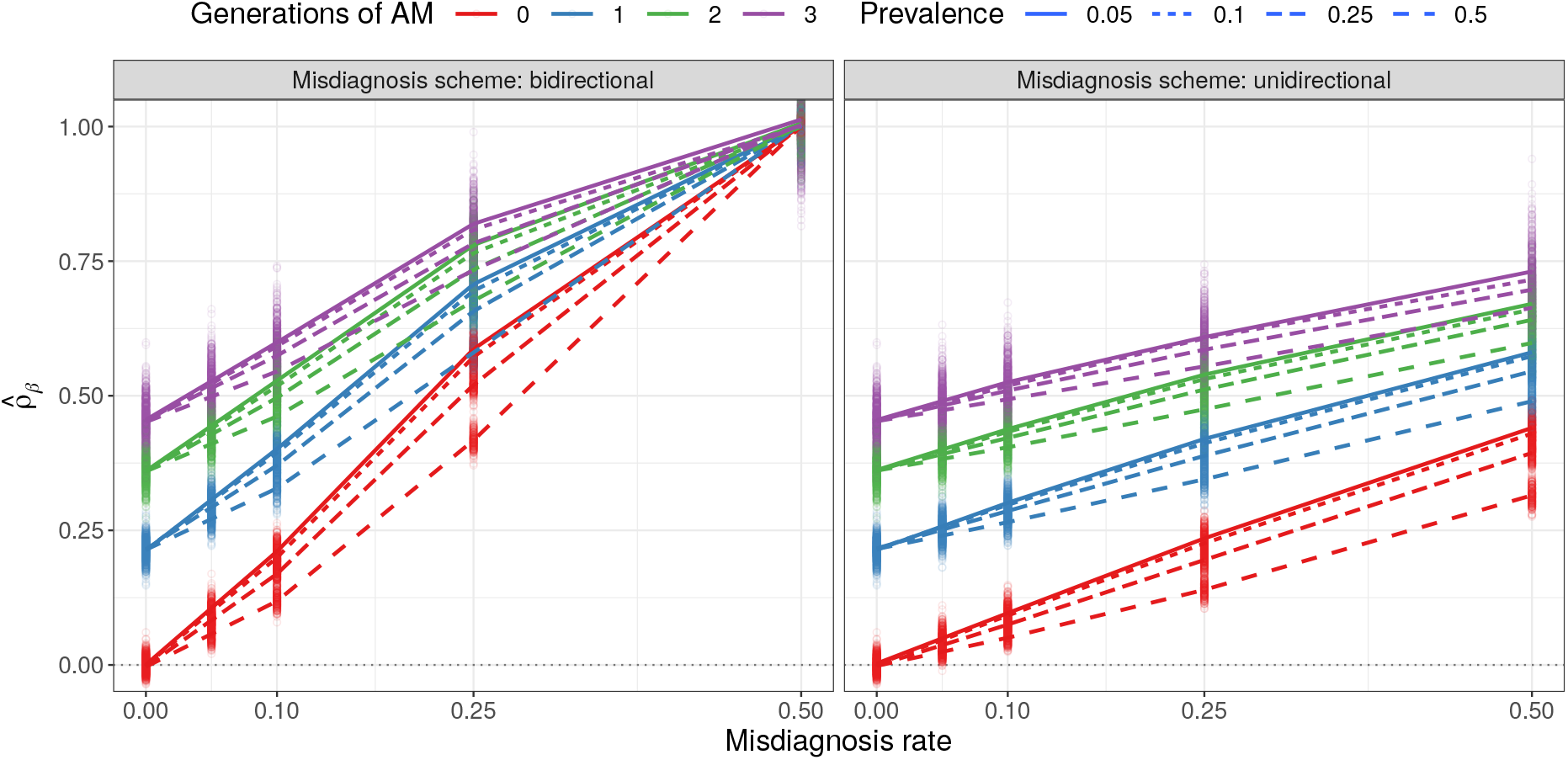
Impact of xAM and misdiagnosis errors on effect correlation estimates between genetically orthogonal binary traits. Panmictic heritabilities and cross-mate correlations are all fixed at 0.5. Under the unidirectional scheme, individuals with disorder *A* are mislabeled as controls for disorder *A* and cases for disorder *B*, regardless of their true status for disorder *B*, at the rate reflected on the *x* axis. Under the bidirectional scheme, the analogous misdiagnoses are enforced for disorder *B* as well. The induced bias in both cases in more pronounced for less common disorders.

**Figure S7:**
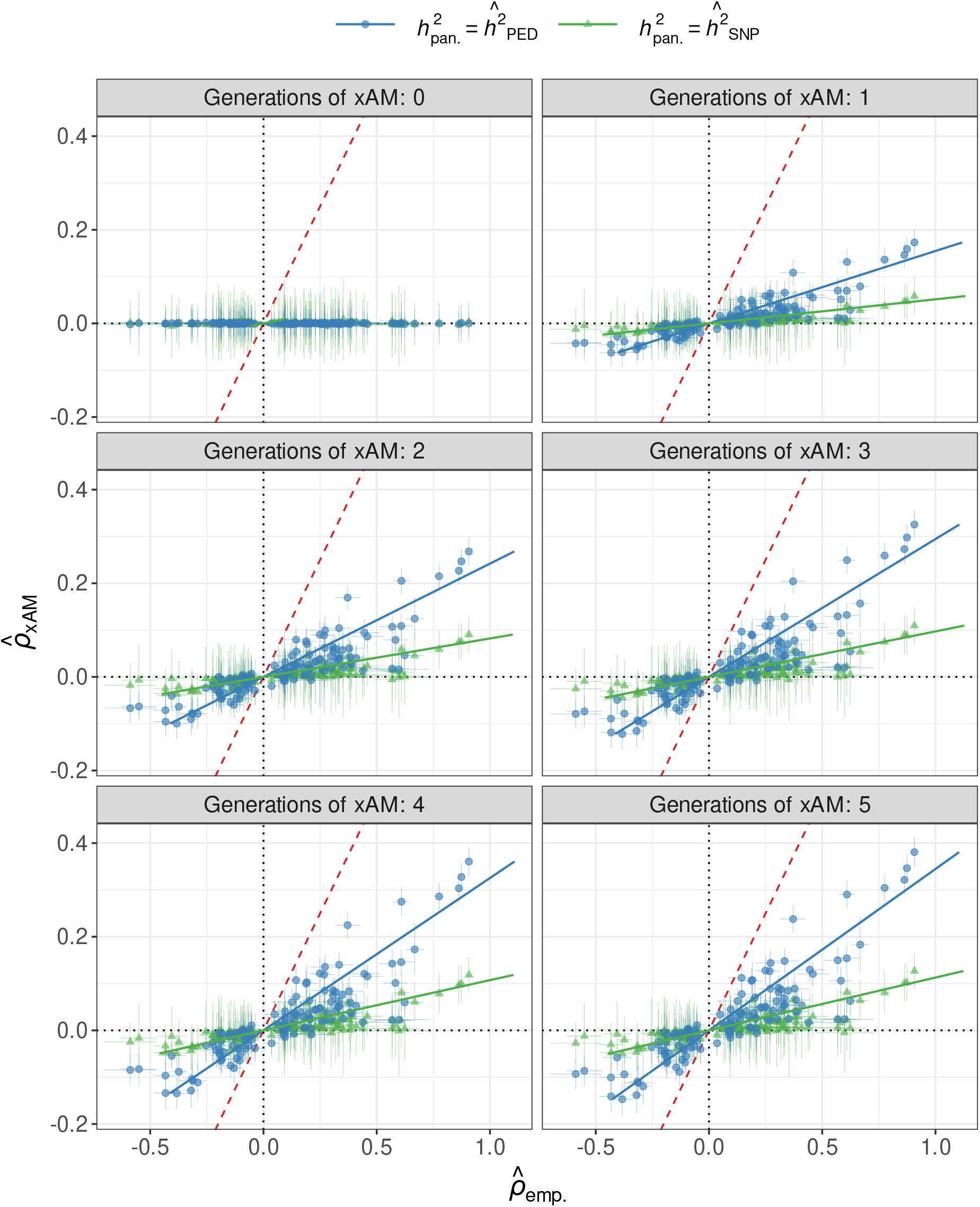
Projected versus empirical 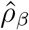 for UK Biobank trait pairs as a function of number of generations of xAM.

**Figure S8:**
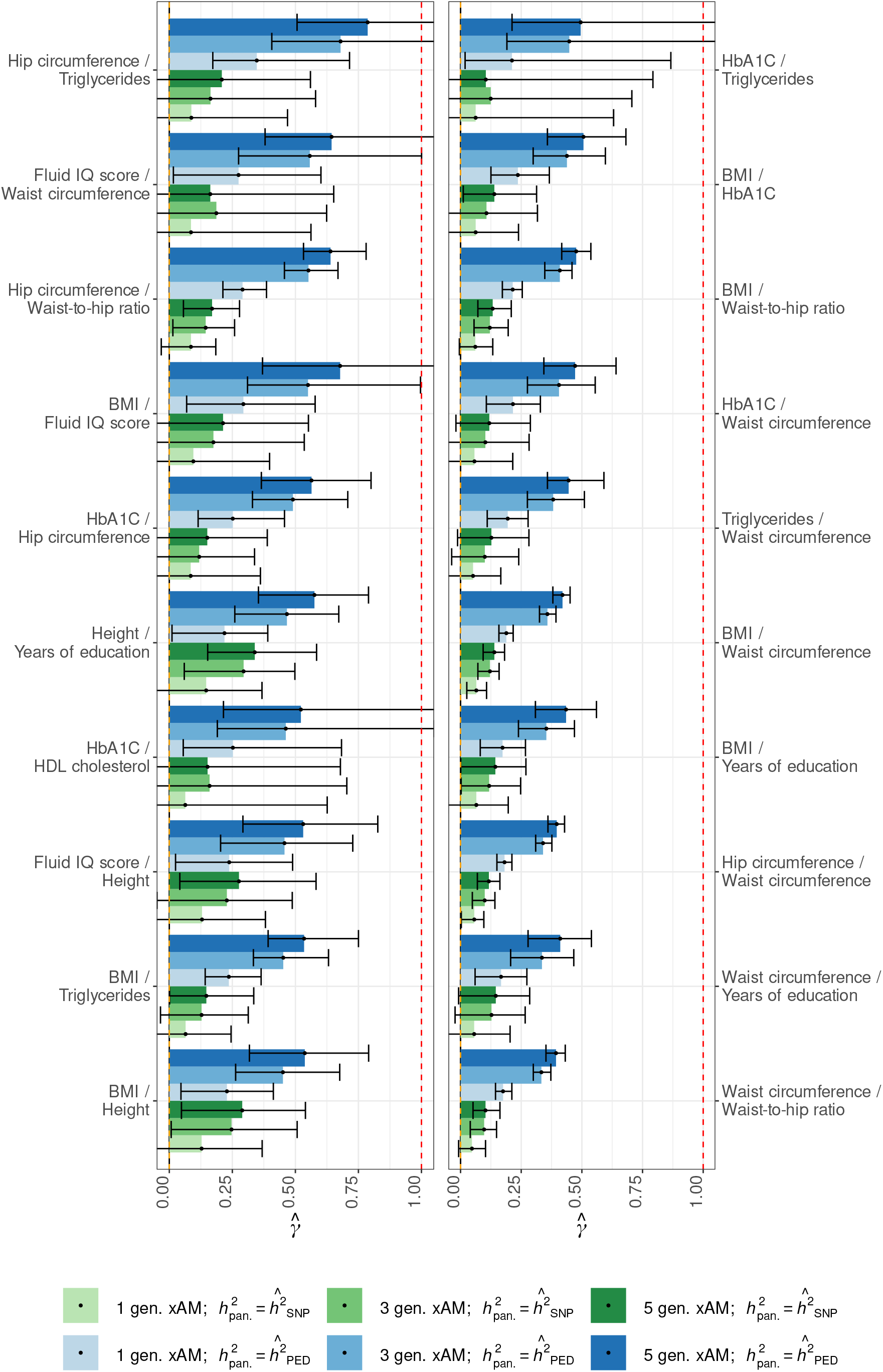
Projected 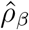 estimates relative to empirical estimates for UK Biobank trait pairs (1 of 2).

**Figure S9:**
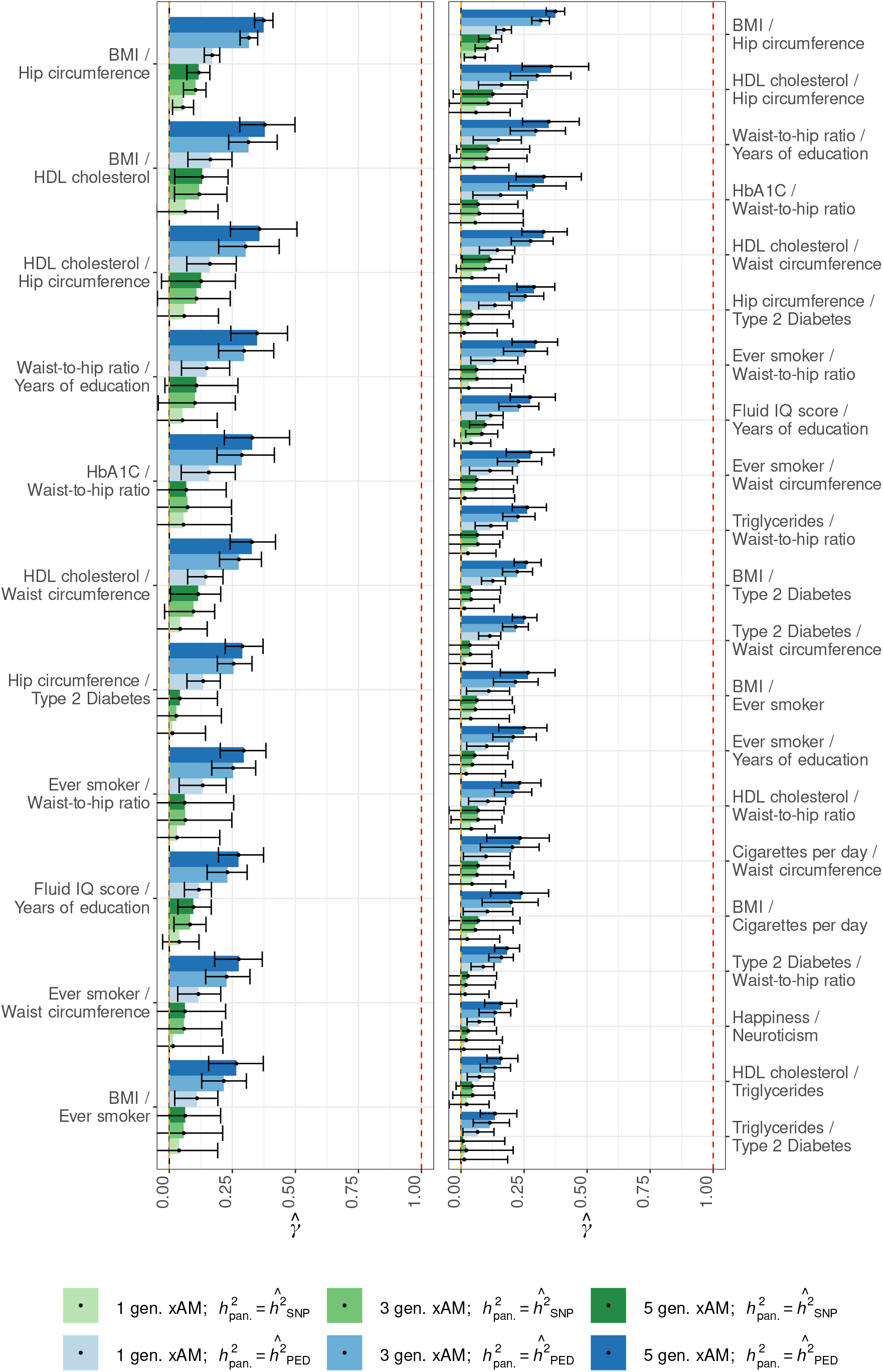
Projected 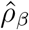 estimates relative to empirical estimates for UK Biobank trait pairs (2 of 2).

**Figure S10:**
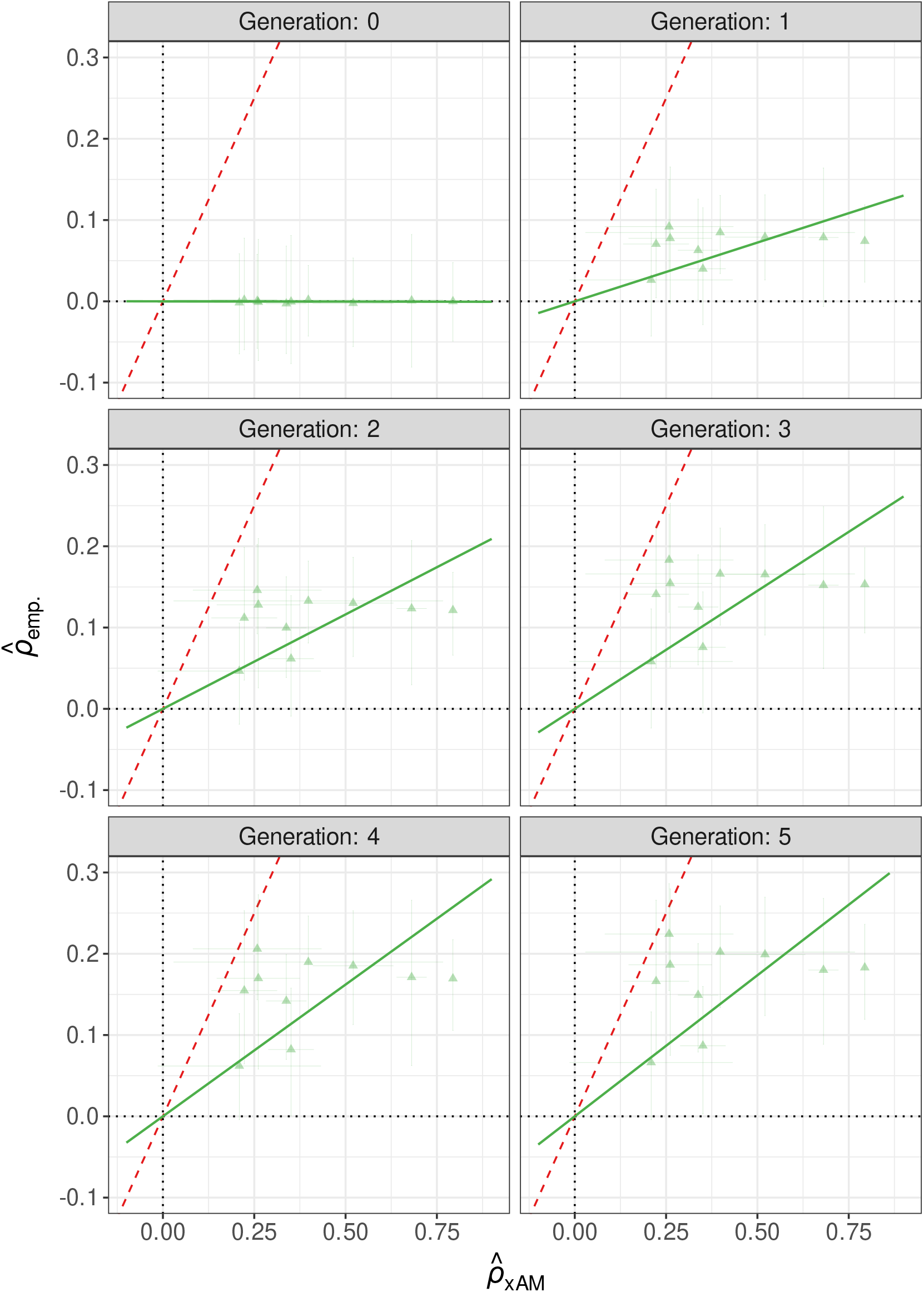
Projected versus empirical 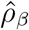 for psychiatric trait pairs

**Figure S11:**
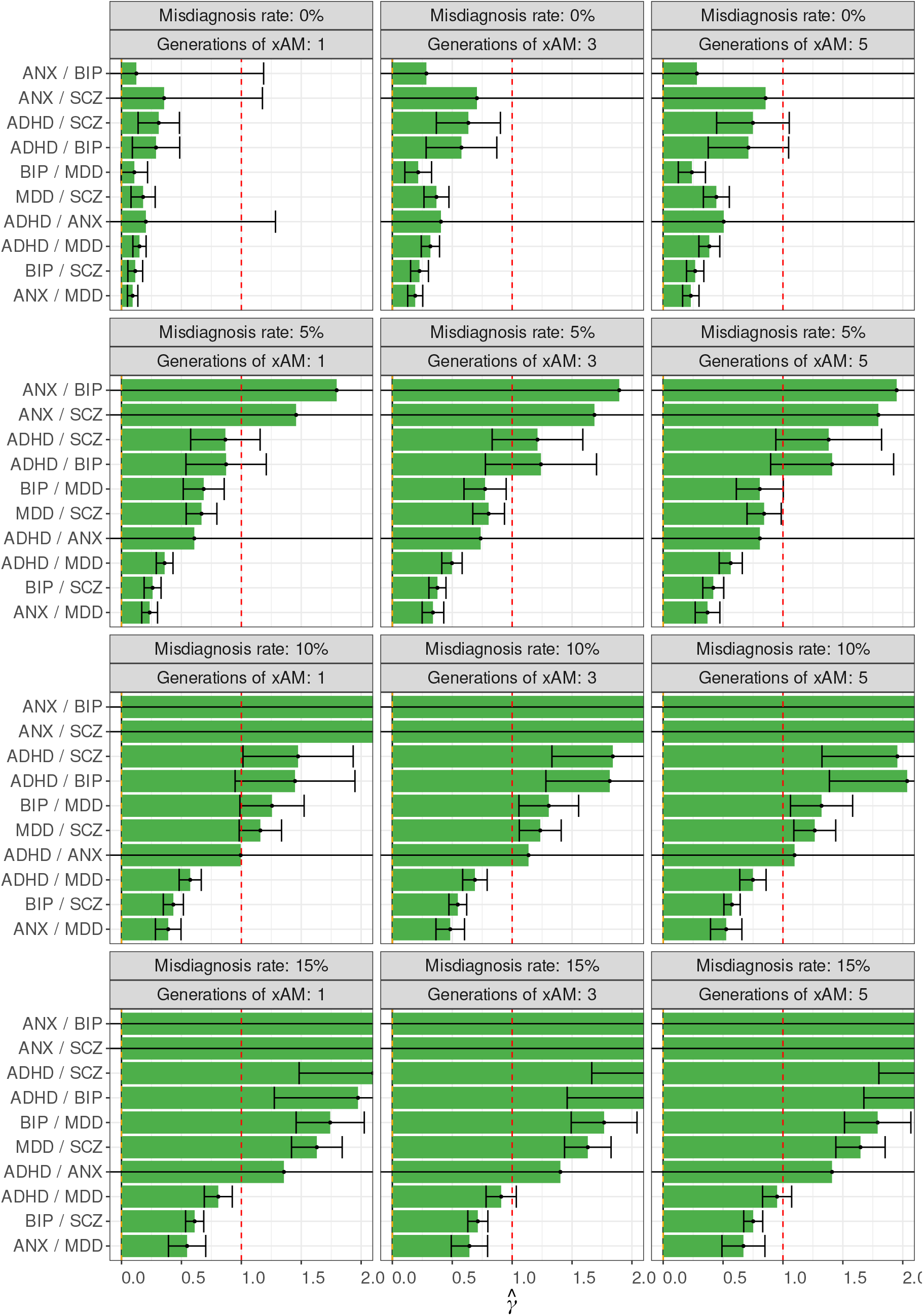
Projected 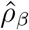 estimates relative to empirical estimates for psychiatric trait pairs under xAM and misdiagnosis rates

**Figure S12:**
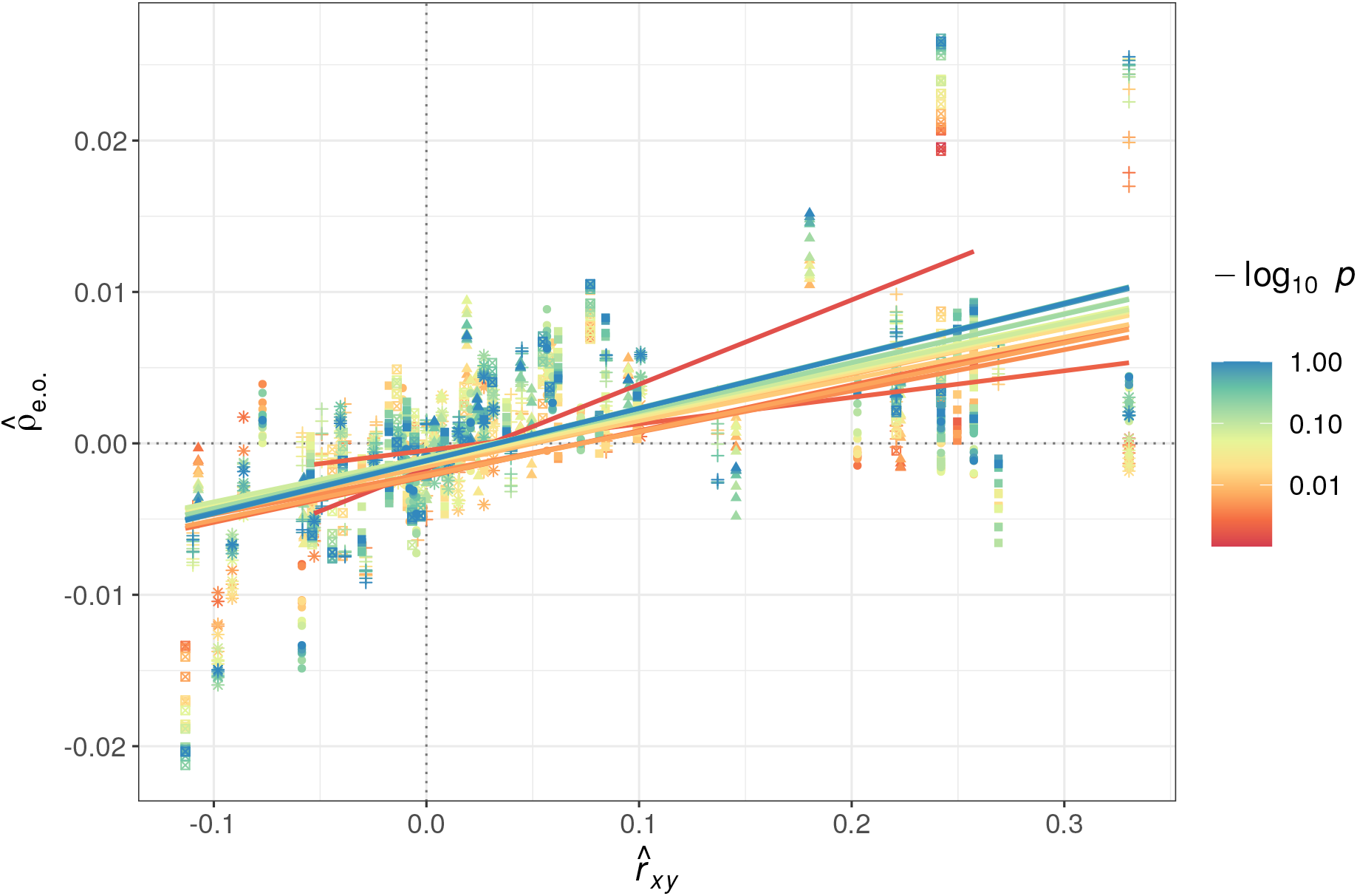
Cross-chromosome polygeneic score correlations and cross-mate phenotypic correlations for varying *p*-value thresholds

